# Plant-specific adaptations of the CDC48 unfoldase

**DOI:** 10.1101/2025.03.28.645892

**Authors:** Brandon Huntington, Anandsukeerthi Sandholu, Jun Wang, Junrui Zhang, Lingyun Zhao, Bilal M. Qureshi, Umar F. Shahul Hameed, Stefan T. Arold

**Author notes:** Correspondence: UFSH, STA.

## Abstract

Targeted protein degradation through the CDC48 unfoldase enables the maintenance and rapid adaptation of proteomes across eukaryotes. However, the profound differences between animals, fungi, and plants are expected to have led to a significant adaptation of the CDC48-mediated degradation. While animal and fungal CDC48 systems have shown structural and functional preservation, such analysis is lacking for plants. We determined the structural and functional characteristics of *Arabidopsis thaliana* CDC48A in various states and bound to the target-identifying cofactors UFD1 and NPL4. Our analysis reveals several features that distinguish *At*CDC48 from its animal and yeast counterparts, despite an 80% sequence identity. Key features are that *At*CDC48A displays distinct domain dynamics and interacts differently with *At*NPL4. Moreover, *At*NPL4 and *At*UFD1 do not form an obligate heterodimer, but independently bind to *At*CDC48A and mediate target degradation; however, their joint action is synergistic. An evolutionary analysis supports that these *Arabidopsis* features are conserved across plants and represent the ancestral state of eukaryotic CDC48 systems. Jointly, our findings support that plant CDC48 retains a greater modular and combinatorial cofactor usage, highlighting a specific adaptation of targeted protein degradation in plants.

## INTRODUCTION

Targeted protein degradation is essential for maintaining a healthy proteome and enabling organisms to adapt to changing environments and metabolic demands (Harper and Bennett, 2016). The segregase and unfoldase, cell division cycle protein 48 (CDC48), is at the heart of this function by coupling the chemical energy of ATP hydrolysis into mechanical force, facilitating protein extraction, unfolding, and targeting to proteasomal degradation (Bodnar and Rapoport, 2017b; Inès et al., 2024).

Studies using purified complexes have provided insights into the structure and function of CDC48 in animals (*Homo sapiens*, *Hs*p97) and yeast (*Saccharomyces Cerevisiae, Sc*Cdc48) (Banerjee et al., 2016; Cooney et al., 2019; Twomey et al., 2019; Pan et al., 2021a; Pan et al., 2021b). CDC48, a AAA+ ATPase, assembles into a homo-hexameric barrel comprising two stacked rings with a central pore. Each protomer includes a flexible N-terminal extension, an N-terminal domain (NTD), two sequential ATPase domains (D1 and D2), and a C-terminal extension (**Figure 1A**). The NTD and C-terminal extension recruit pathway-specific cofactors (Buchberger et al., 2015, 97; Blueggel et al., 2023) and undergo regulatory post-translational modifications (Madeo et al., 1998; Hänzelmann and Schindelin, 2017). D1 and D2 facilitate ATP hydrolysis and generate the mechanical force required for substrate unfolding via flexible pore loops in each domain (Pan et al., 2021b; Ji et al., 2022).

**Figure 1.**
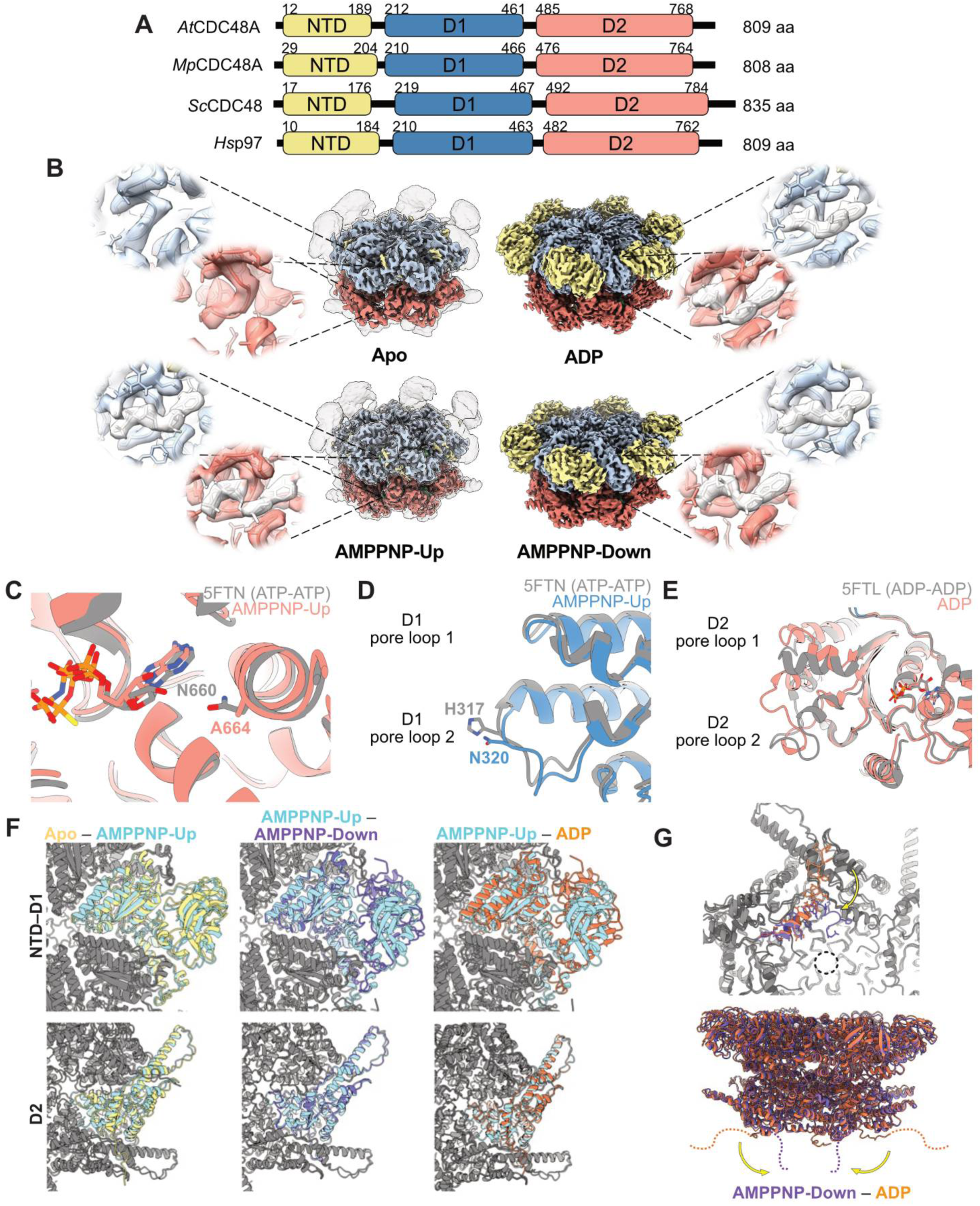
Idiosyncrasies of *At*CDC48A. **A)** Domain architecture comparison of CDC48 in *Arabidopsis thaliana* (*At*), *Marchantia polymorpha* (*Mp*), *Saccharomyces cerevisiae* (*Sc*), and *Homo sapiens* (*Hs*). The N-terminal domain (NTD) is coloured yellow. The D1 and D2 ATPase domains are coloured blue and red, respectively. Amino acid (aa) numbers are provided for each domain. **B)** Cryo-EM reconstructions of *At*CDC48A nucleotide states (apo: **EMD-63608**, AMPPNP-Up: **EMD-63609**, AMPPNP-Down: **EMD-63610**, ADP: **EMD-63611**). The unsharpened maps are shown as grey transparent surfaces over the sharpened maps of apo and AMPPNP-Up *At*CDC48A to highlight the flexible regions. Domains are coloured according to Panel A, and the nucleotides are coloured white. Spotlights show the D1 and D2 nucleotide binding pockets with atomic models overlaid (apo: PDB **9M3V**, AMPPNP-Up: PDB **9M3W**, AMPPNP-Down: PDB **9M3X**, ADP: PDB **9M3Y**). **C-E)** Comparison of *At*CDC48A and *Hs*p97. **C)** Comparison of the D2 active site. *At*CDC48A (PDB **9M3W**) is coloured red, *Hs*p97 (PDB **5FTN**) is coloured grey. AMPPNP and A664 (*At*CDC48A) and ATPᵧS and N660 (*Hs*p97) are represented as sticks. **D)** Comparison of D1 pore loops. AMPPNP-Up *At*CDC48A (PDB **9M3W**) is coloured blue, ATPᵧS-bound *Hs*p97 (PDB **5FTN**) is coloured grey. **E)** Comparison of D2 pore loops. ADP-bound *At*CDC48A (PDB **9M3Y**) is coloured blue, ADP-bound *Hs*p97 (PDB **5FTN**) is coloured grey. ADP molecules are represented as sticks. **F)** Comparison of the NTD and D1 (top) and D2 (bottom) between the apo and AMPPNP-Up states (left), the AMPPNP-Up and AMPPNP-Down states (middle) and AMPPNP-Up and ADP states (right). Apo *At*CDC48A is coloured yellow, AMPPNP-Up is coloured blue, AMPPNP-Down is coloured purple, and ADP-bound *At*CDC48A is coloured purple. Only one protomer is coloured for clarity. **G)** Bottom (top) and side view (bottom) of the C-terminal extension positioning in AMPPNP-Down and ADP-bound *At*CDC48A. Black dotted lines (top) represent the pore of the hexamer. Coloured dashed lines (bottom) represent assumed continuation of the C-terminal extension. Coloured as in Panel F.

CDC48 interacts with its targets through two classes of adaptor proteins. The first class includes the ubiquitin fusion degradation protein 1 (UFD1) and nuclear protein localisation protein 4 (NPL4) system, observed as an obligate heterocomplex in humans and yeast, which recruits polyubiquitinated substrates(Sato et al., 2019; Nguyen et al., 2022). The second class comprises the large and diverse family of ubiquitin regulatory X (UBX)–containing proteins. UBX-family cofactors have evolved diverse functions beyond the recruitment of ubiquitinated substrates, including recognising non-ubiquitinated substrates, or the catalytic control of CDC48 through hexamer disassembly or recruitment of inactivating factors (Rancour et al., 2004; Alexandru et al., 2008; Marshall et al., 2019; Kracht et al., 2020, 20020; Zhang et al., 2021; Zhang et al., 2022). Target-binding cofactors bring the substrate in proximity to CDC48 by interacting with its NTD and/or D1. The substrate is partially unfolded, and the unstructured region is inserted into the pore of the CDC48 hexamer, where D2 pore loops mediate translocation, possibly by a hand-over-hand mechanism common to other AAA+ ATPases (Twomey et al., 2019; van den Boom et al., 2023). D2 primarily drives translocation due to its strongly interacting aromatic pore-loop residues, whereas D1 ATPase activity regulates substrate release alongside a deubiquitinase processing cofactor (Bodnar and Rapoport, 2017a; Bodnar and Rapoport, 2017b).

Several mechanistic aspects remain unclear, including the regulation of CDC48 interactions with cofactors and targets, the existence of additional cofactor pathways that initiate unfolding, the conservation of unfolding mechanisms across species, and the process of substrate transfer from CDC48 to the proteasome.

Rapid proteomic changes mediated by CDC48 are particularly critical in sessile plants to enable them to adapt to environmental changes. Plants also differ fundamentally from yeast and animals in their cellular and organismal structures, suggesting that their CDC48 system has undergone substantial modifications. Yet, despite approximately one billion years of evolutionary divergence, plant CDC48 orthologues share nearly 80% sequence identity with their animal and fungal counterparts. This conservation raises questions about how this system has adapted to plant-specific roles, such as growth, germination, flowering, chloroplast regulation, and immunity (Rancour et al., 2002; Park et al., 2008; Copeland et al., 2016; Bègue et al., 2019; Ao et al., 2021; Rosnoblet et al., 2021; Li et al., 2022; Gao et al., 2022b; Inès et al., 2024). However, detailed structure–function analyses of plant CDC48s are currently lacking.

Our study reveals the functional and structural characteristics of *Arabidopsis thaliana* CDC48A (*At*CDC48A), alone and bound to the main adaptor proteins *At*NPL4 and *At*UFD1. We observe subtle differences in the structure and dynamics of *At*CDC48A compared to its human and yeast orthologues. However, pronounced differences are observed in the way that *At*CDC48 connects with *At*NPL4, and *At*UFD1, where plants have retained a more ancestral independent usage of the adaptors. Collectively, these findings provide insights into the evolution and adaptation of CDC48-based protein unfolding and degradation across eukaryotic lineages.

## RESULTS

### Idiosyncrasies of Arabidopsis CDC48A compared to animal and fungal orthologues

The domain structure and sequence of CDC48 are highly conserved across higher and lower plants and across eukaryotic kingdoms, with *At*CDC48A sharing 88% sequence identity with the liverwort *Marchantia polymorpha* (*Mp*CDC48A) and 77% and 66% sequence identity with *Hs*p97 and *Sc*Cdc48, respectively (**Figure 1A**; **Supplemental Figure 1**). To identify plant-specific adaptations, we collected single-particle cryo-EM datasets of recombinant full-length *At*CDC48A, using a previously described method to mitigate preferential orientation by targeting regions of thicker ice (Huntington et al., 2022). Four structures of *At*CDC48A were determined: ADP-bound (3.0 Å resolution), two ATP analogue (AMPPNP) states (AMPPNP-Up and AMPPNP-Down; 3.2 Å and 3.3 Å resolution), and the apo-state without nucleotide (3.9 Å resolution) (**Figure 1B; Supplemental Table 1; Supplemental Figures 2–10**). Additionally, we determined the crystal structure of ADP-bound *At*CDC48A (residues 28-809) to 3.4 Å, and the 2.3 Å crystal structure of the unliganded NTD (residues 28-190) (**Supplemental Table 2**). For all structures, the nature or absence of nucleotides was visible in the density maps (**Figure 1B**).

Consistent with its high sequence conservation, *At*CDC48A shares several structural similarities with animal and fungal orthologues: (i) The catalytic D1 and D2 domains of *At*CDC48A include conserved active site motifs, including the Walker A (K254/K527, T255/T528), Walker B (D307/D580, E308/E581), Sensor 1 residues (N351/N628), the sensor loop, and the short region of homology (SRH) featuring arginine fingers (R362, R365/R639, R642, R770) (**Supplemental Figure 1**). (ii) In the ADP-bound state, the NTD aligned coplanar with the D1 ring and is stabilised by contacts with D1. Conversely, in the AMPPNP-bound and apo-states, the NTD adopted a more flexible position above the D1 ring, resulting in poorly resolved regions in cryo-EM density maps across structures (**Figure 1B**). (iii) The NTD conformation remains largely consistent between D1-bound states and the crystal structure of the isolated NTD (RMSD 1.0 Å - 1.5 Å), except for a loop region (residues 120 - 128), which was unresolved in the crystal structure but visible in the AMPPNP-Down and ADP-bound cryo-EM data, where it was stabilised by D1 interactions (**Supplemental Figures 11 and 12**).

We identified several differences between *At*CDC48A and its animal or fungal orthologues. First, we observed a subtle difference in the D2 active site, where *At*CDC48A featured an alanine (A664) instead of an asparagine (N660 in *Hs*p97), which interacts with the ribose/adenine moiety. This substitution may influence nucleotide binding and positioning, potentially affecting ATP hydrolysis rates in plants (**Figure 1C**). Second, in the D1 pore loop 2, *At*CDC48A substituted the histidine gate present in *Hs*p97 (H317) for asparagine (N320) which may influence substrate processing rates by altering zinc ion coordination (DeLaBarre and Brunger, 2003) or π-π stacking interactions with the D2 pore loop tryptophan (W551) (Pan et al., 2021b) (**Figure 1D**). Third, AMPPNP-bound *At*CDC48A clearly exhibited an NTD-down position, rarely reported in cryo-EM structures of orthologues (Schuller et al., 2016; Gao et al., 2022a). Finally, we only observed single hexamers in cryo-EM *At*CDC48A datasets, and a NTD-D1–mediated hexamer–hexamer stacking in the crystallographic *At*CDC48A structure (**Supplemental Figure 12B**), whereas *Hs*p97 hexamers can also form a D2-mediated dodecameric state, particularly in nucleotide-free conditions (Hoq et al., 2021; Nandi et al., 2021; Caffrey et al., 2021; Gao et al., 2022a; Arie et al., 2024; Nandi et al., 2024).

### Structural dynamics of *At*CDC48A

Comparison of *At*CDC48A’s catalytic domains across AMPPNP, ADP, and apo states revealed ∼1 Å shifts in the active sites of the apo-state relative to nucleotide-bound states, with minimal differences between AMPPNP-Up and AMPPNP-Down conformations. In the ADP state, D1 pore loop 1 was 2.5 Å wider, while the D1 pore loop 2 was 2 Å narrower than in the AMPPNP states, and D2 pore loop 2 density was resolved more clearly (**Figure 1E**). The D2 pore loops in the ADP state also resembled the AMPPNP-bound *At*CDC48A conformation more closely than ADP-bound *Hs*p97 (**Figure 1E**). Unlike *Hs*p97, no substantial rotation of D2 relative to D1 was observed in different nucleotide states, with the pore size also remaining consistent as measured by the position of the terminal helix ⍺9 (**Figure 1F**) (Noi et al., 2013; Banerjee et al., 2016). Additionally, in nucleotide-free or ADP-bound states, the C-terminal extension 3F repeat (residues F772–F778) interacted with the hydrophobic cleft between ⍺6 and ⍺8 of the adjacent protomer. In contrast, in AMPPNP-bound states, the C-terminal extension contributed an arginine finger (R770) to the D2 active site, reducing interprotomer interactions and priming the complex for structural changes necessary during substrate processing (**Figure 1G**).

To examine the dynamic features of *At*CDC48A, we analysed the cryo-EM datasets using 3D variability analysis (3DVA) in CryoSPARC. The ADP-bound state exhibited the lowest heterogeneity, while the apo-state showed moderate heterogeneity and the AMPPNP-bound state displayed high heterogeneity, particularly in NTD positioning (**Figure 2**). Analysing two orthogonal eigenvectors of the AMPPNP-bound state produced five reconstructions each, discretizing a continuum of states. Along the first eigenvector, the six NTDs exhibited continuous heterogeneity, transitioning between AMPPNP-Up and AMPPNP-Down conformations (**Figure 2A; Movie 1**). Along the second eigenvector, three NTDs were positioned up while the other half were positioned down (**Figure 2B; Movie 2**). In contrast, variability analysis of the apo-state did not suggest a similar extent of coordinated NTD motion, instead revealing flexibility within the enzyme core, likely due to reduced inter-protomer interactions in the absence of bound nucleotides (**Movie 3**).

**Figure 2:**
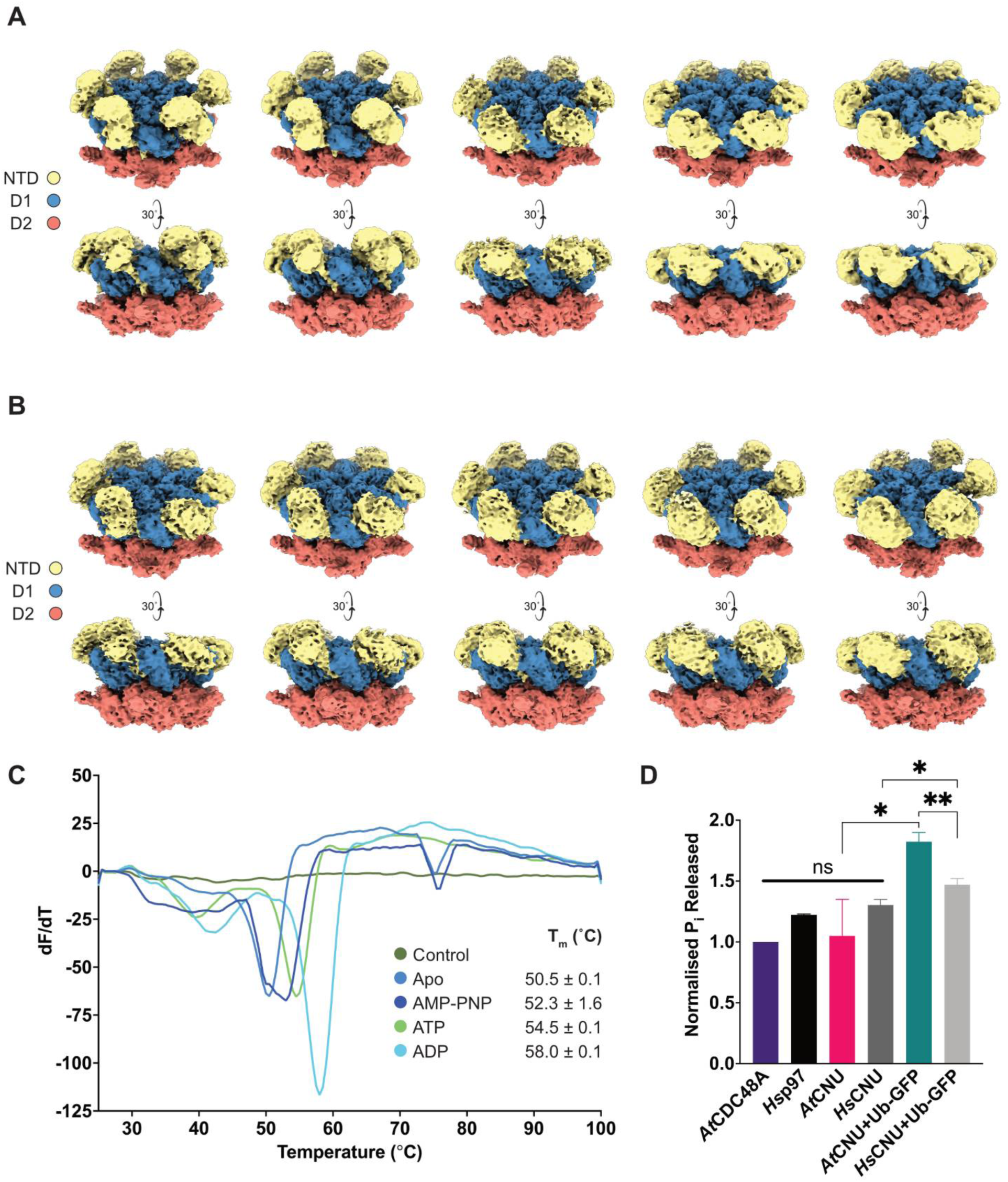
Structural dynamics of *At*CDC48A. **A)** Cryo-EM reconstructions clustered along the first 3DVA eigenvector of AMPPNP-bound *At*CDC48A highlighting the movement of all six NTDs together. Domains are coloured as in Figure 1. **B)** Cryo-EM reconstructions clustered along the second 3DVA eigenvector of AMPPNP-bound *At*CDC48A highlighting the movement of half of the NTDs together. Domains are coloured as in Figure 1.**C)** Differential scanning fluorimetry of *At*CDC48A in different nucleotide-bound states. Rate of change of fluorescence with temperature for apo (dark blue) AMPPNP-bound (purple), ATP-bound (green) and ADP-bound (light blue) *At*CDC48A was measured, using the Sypro Orange dye alone as control (dark green). The melting temperature (T_m_) is reported as mean ± standard deviation (n=3). **D)** ATPase activity of *A. thaliana* (*At*) and *H. sapiens* (*Hs*) CDC48 (purple and black, respectively) and CNU complexes (pink and dark grey, respectively) alone or with ubiquitinated GFP (green and light grey, respectively). For each replicate, the fluorescence intensity was normalised to the value of *At*CDC48A alone. Normalised fluorescence values are reported as mean ± SEM (n=3). A one-way ANOVA found no significance (ns) between *At*CDC48A and substrate-free complexes (p > 0.05). Stars represent significant differences between pairs, calculated by unpaired t-test (* represents p < 0.05; ** represents p < 0.01). Statistical tests were performed using Prism (v10.0).

Differential scanning fluorimetry (DSF) experiments correlated with the relative stability of the *At*CDC48A complex across various nucleotide conditions. The nucleotide-free *At*CDC48A was the least heat stable compared to AMPPNP and ATP states, which, in turn, were less stable than the ADP-bound *At*CDC48A (melting temperatures, *T_m_*, were 50.5 ± 0.1, 52.3 ± 1.6, 54.5 ± 0.1 and 58 ± 0.1 °C, respectively) (**Figure 2C**). Additionally, ATP hydrolysis assays showed *At*CDC48A had a basal ATP turnover rate comparable to its human orthologue, *Hs*p97, in the absence of substrate (**Figure 2D**).

Combined, our analysis of different nucleotide states provides insights into the catalytic cycle of *At*CDC48A. When nucleotide-free, *At*CDC48A displays a significant level of malleability, and the NTDs are positioned flexibly above the plane of the D1 ring. The binding of ATP stabilises *At*CDC48A overall but also enables the NTDs to oscillate between an ‘Up’ and ‘Down’ conformation, priming the complex for cofactor engagement and subsequent hydrolysis to ADP. ATP binding at D1 enables the N-terminal extension to form a 3_10_ helix and stably interact with the neighbouring D1 when the NTDs are positioned Up. ATP binding at D2 induces disorder in the pore loops preparing them for substrate interaction, while changes in D2 ⍺6 and ⍺8 allow the C-terminal extension to detach from the neighbouring protomer and contribute an arginine finger to the neighbouring D2 active site. Upon hydrolysis to ADP, the NTD stabilises on the D1 domain, causing shifts in the D1 pore loops, and no large-scale rotation of D2 occurs. The D2 pore loops transition from disorder to order, facilitating substrate translocation, and the C-terminal extension reintegrates on the neighbouring protomer to further stabilise the moved complex. Although most features were conserved across kingdoms, the NTD dynamics and lack of D2 rotation may imply differences in substrate processing of plants compared to orthologs.

### Plant-specific connections between CDC48, UFD1 and NPL4

Across kingdoms UFD1 preserves its overall architecture consisting of an N-terminal UT3 domain (which is structurally similar to the CDC48 NTD (Pye et al., 2007)), and a C-terminal flexible region containing linear interaction motifs. Overall, both paralogues of *At*UFD1 (AtUFD1A and *At*UFD1B) share 65% sequence identity with *Mp*UFD1 and 40-50% with *Hs*Ufd1 or *Sc*Ufd1, with the UT3 domain being more conserved than the C-terminal flexible regions (78% sequence identity with *Mp*UFD1 and 50-60% with *Hs*Ufd1 or *Sc*Ufd1) (**Figure 3A**). In humans and yeast, UFD1 contains two NTD-binding SHP motifs (SHP1 and SHP2), characterised by the consensus sequence *FxGxGxxL* and an NPL4 binding motif (NBM) with the consensus sequence *hxhxhGxhxh*, where x represents any amino acid and *h* denotes hydrophobic residues (**Figure 3B**). Whereas SHP1 is conserved across species, the second glycine of SHP2 is replaced by a lysine in *At*UFD1 and *Mp*UFD1 (**Figure 3B**). Additionally, *At*UFD1 and *Mp*UFD1 lack the second and third hydrophobic residues in the NBM, and the first hydrophobic residue is less prominent (alanine or proline instead of leucine or tyrosine) (**Figure 3B**). AlphaFold modelling supported an interaction between CDC48–UFD1, but not NPL4–UFD1, in *Arabidopsis* and *Marchantia* (**Figure 3C**; **Supplemental Figures 13 and 14**). These features suggested that the interaction between the UFD1 SHP1 motif and CDC48 is conserved in plants, but the interaction between UFD1 and NPL4 is likely disrupted due to alterations in the NBM.

**Figure 3:**
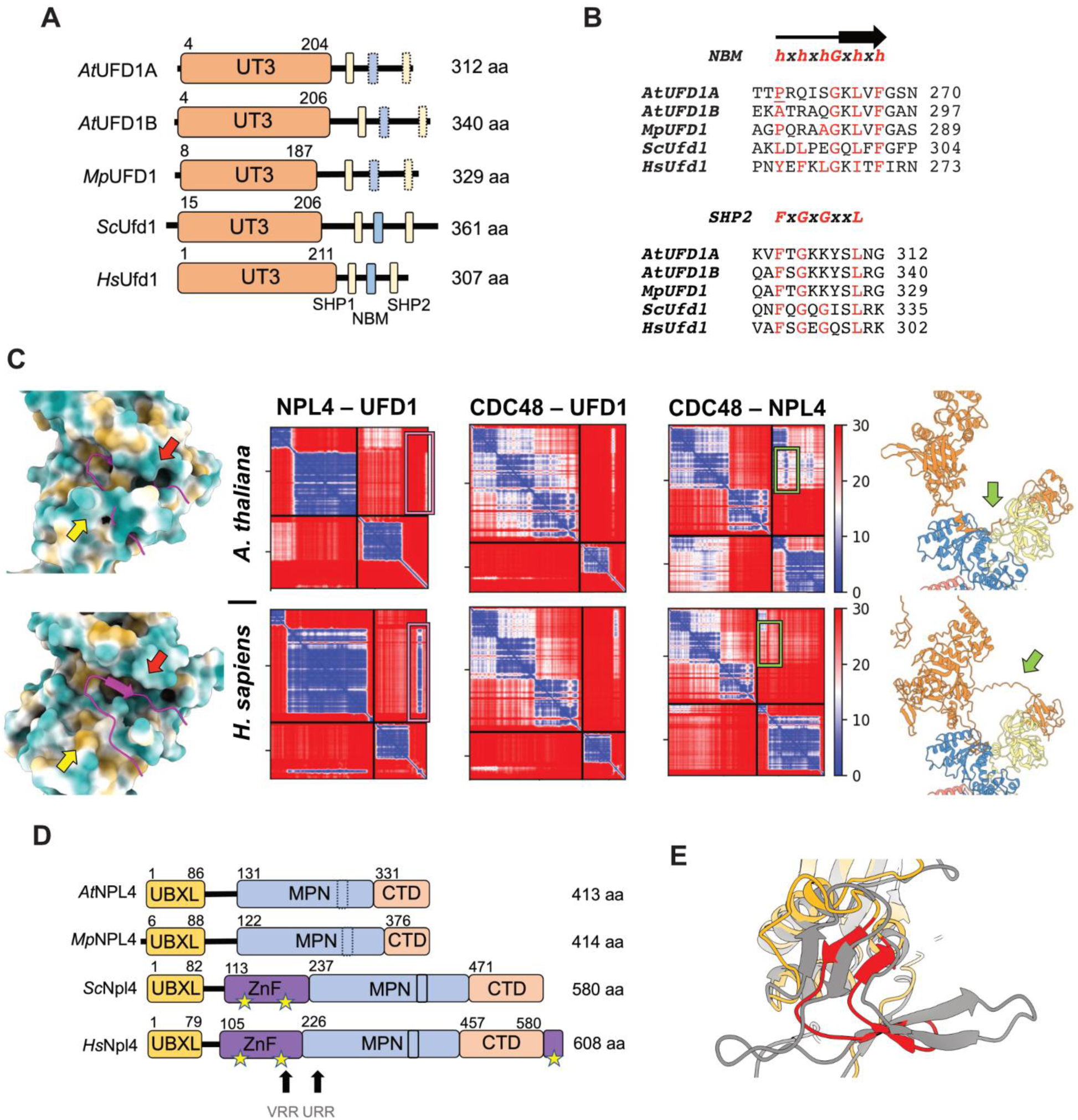
Domain and predicted interaction differences of *At*NPL4 and *At*UFD1. Species names are abbreviated as: *Arabidopsis thaliana* (*At*), *Marchantia polymorpha* (*Mp*), *Saccharomyces cerevisiae* (*Sc*), and *Homo sapiens* (*Hs*). **A)** Domain architecture comparison of UFD1 across species. Boxes indicate binding motifs and dashed lines indicate motif differences between orthologues. The UT3 domain is coloured orange. The SHP1 and SHP2 motifs are coloured yellow. The NPL4-binding motif (NBM) is coloured blue. Amino acid (aa) numbers are provided for each domain. **B)** Sequence alignment of the UFD1 NBM (top) and SHP2 (bottom) motifs. The consensus sequence is provided above the sequence alignment. A cartoon representation of the secondary structure of the NBM is provided above the consensus sequence. Conserved residues are coloured red. **C)** Domain architecture comparison of NPL4. The UBXL domain is coloured orange. The zinc-finger (ZnF) domains are coloured purple. The MPN domain is coloured blue. The C-terminal domain (CTD) is coloured peach. Boxes indicate binding motifs and dashed lines indicate motif differences between orthologues. Stars represent canonical ZnF motifs. Arrows highlight the *Hs*p97/VCP-regulatory region (VRR) and *Hs*UFD1-regulatory region (URR) of *Hs*Npl4. Amino acid (aa) numbers are provided for each domain. **D)** AlphaFold predicted structures of the insert-2 region from the MPN domain of *At*NPL4 (coloured) and *Hs*Npl4 (grey). The insert-2 region of *At*NPL4 is coloured red. **E)** Pairwise predicted interactions of *A. thaliana* (top) and *H. sapiens* (bottom) CNU proteins using AlphaFold. Predicted aligned error (PAE) plots for the predicted model of UFD1–NPL4 (left), CDC48–UFD1 (middle) and CDC48–NPL4 (right), shown with scale bar representing low (blue) to high (red) PAE. On the bottom left, a surface representation of *Hs*Npl4’s UFD1-interacting region is depicted (coloured blue–gold for hydrophilic–hydrophobic residues) overlaid with cartoon representation of *Hs*Ufd1’s NBM (pink), corresponding to the NPL4–UFD1 PAE plot (pink box, left). On the top left, *Hs*Ufd1 is overlaid on *At*NPL4 to show the binding site occlusion. Arrows represent the UFD1 interaction sides on NPL4. Red arrows indicate the cleft of β-sheet formation and yellow arrows represent the hydrophobic patch. The predicted models of *At*CDC48A–*At*NPL4 (top right) and *Hs*p97– *Hs*Npl4 (bottom right) with green arrows showing NPL4’s UBXL–MPN linker region, corresponding to the CDC48–NPL4 PAE plot (green box, right). The CDC48 NTD, D1, and D2 are coloured yellow, blue, and red, respectively. NPL4 is coloured orange.

The Mpr/Pad1 N-terminal (MPN) domain of NPL4 is responsible for binding UFD1 and stabilising unfolded ubiquitin (Twomey et al., 2019; Sato et al., 2019; Nguyen et al., 2022) (**Figure 3D**). The UFD1-binding region (UBR) of NPL4 consists of a β-strand pocket at the MPN insert-1 region and a hydrophobic patch (Sato et al., 2019; Nguyen et al., 2022). AlphaFold modelling showed that the UBR features were blocked in the MPN domains of *At*NPL4 and *Mp*NPL4, further supporting a disrupted interaction between NPL4 and UFD1 in plants (**Figure 3C; Supplemental Figure 15**). Although *At*NLP4 retains a conserved UBX-like domain (UBXL) which binds to CDC48 NTDs (Bruderer et al., 2004), *At*NLP4 lacks the zinc finger (ZnF) domains that are crucial for CDC48 association in animals and fungi (**Figure 3C**). AlphaFold modelling proposed that the interaction between NPL4 and CDC48 in *Arabidopsis* and *Marchantia* is stabilised through a plant-specific association, involving NPL4 residues of the UBXL–MPN linker, in particular F105 (**Figure 3D; Supplemental Figure 16**). Furthermore, the β-hairpin between β-C and β-D of the MPN insert-2 (also known as the β-strand finger or Npl4 loop) is shorter in plant NPL4 compared to its human and yeast counterparts (**Figure 3E**). Sequence analysis also revealed that plant NPL4 lacks the VCP/p97-regulatory region (VRR), which in *Hs*Npl4 prevents interaction with *Hs*p97 unless *Hs*Npl4 forms a heterodimer with *Hs*Ufd1 (Lass et al., 2008). Thus, although *At*NPL4 interactions and regulatory mechanisms are conserved throughout plants, they differ substantially from those in animals and fungi.

### The Arabidopsis CDC48A–UFD1–NPL4 (CNU) complex shows plant-specific independence of cofactors *in vitro*

We next tested these predictions *in vitro*. All interactions with *At*CDC48A had AMPPNP pre-bound, unless noted. Size-exclusion chromatography coupled with multi-angle light scattering (SEC-MALS) analysis showed that *At*NPL4 with *At*UFD1A or *At*UFD1B did not elute as a complex (**Figure 4A and 4B; Supplemental Figure 17A**). Similarly, microscale thermophoresis (MST) showed no binding between *At*NPL4 and *At*UFD1A at pH 7.5 or 6.5 and no binding between *At*NPL4 and *At*UFD1B at pH 7.5 (K_d_ > 200 µM), and only a thermal shift between *At*NPL4 and *At*UFD1B at pH 6.5 (resulting in an apparent K_d_ of 75.4 +/- 16.5 μM), which was not corroborated by isothermal titration calorimetry (ITC) experiments at this pH, suggesting the MST signal was caused by protein aggregation at high concentrations (**Figure 4C**; **Table 1**; **Table 2**; **Supplemental Figure 17B**). These results confirmed that *At*NPL4 and *At*UFD1 do not form a complex in *Arabidopsis*, conversely to their human and yeast counterparts (Sato et al., 2019; Nguyen et al., 2022).

**Figure 4.**
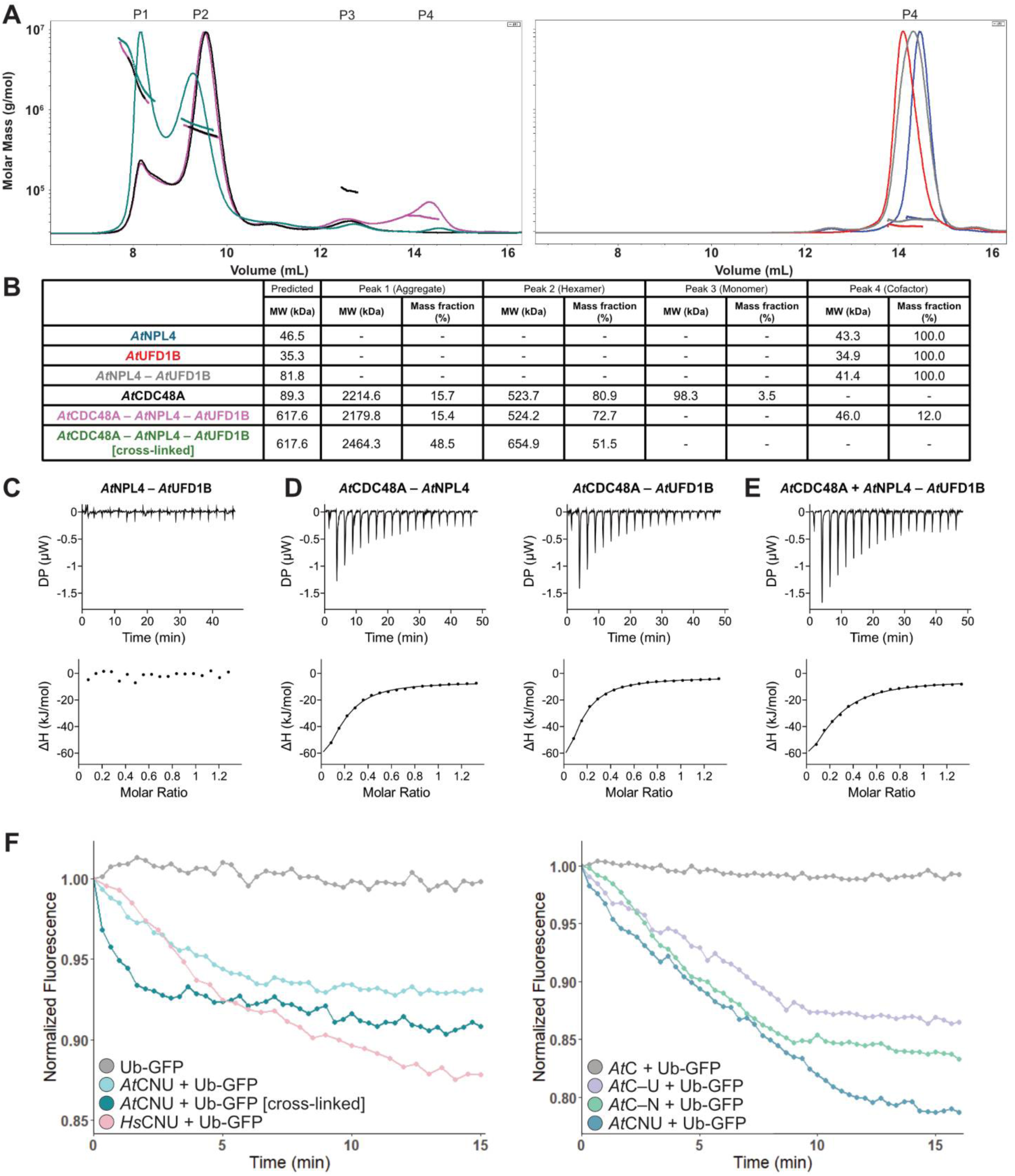
Interaction and function of the *At*CNU complex. **A)** Differential refractive index (solid curves) of *At*CNU proteins measured through Size Exclusion Chromatography coupled with Multi-Angle Light Scattering (SEC-MALS). The calculated molecular weight of each peak (P1–P4) is represented by a dashed line and tabulated in Panel B. For clarity, differential refractive indices are separated by *At*CNU complex comparisons (left) and *At*NPL4–*At*UFD1B comparisons (right). Colours of each species correspond to those in Panel B. **B)** Tabulated molecular weights calculated for each individual *At*CNU protein or complex. The predicted molecular weight of each species is provided, alongside the calculated values for each peak (P1–P4) in Panel A. Colouring is consistent with Panel A, where *At*NPL4 is blue, *At*UFD1B is red, *At*NPL4–*At*UFD1B is grey, *At*CDC48A is black, *At*CNU pink and the crosslinked *At*CNU complex is green. **C-E)** Representative isothermal titration calorimetry (ITC) interaction plots at pH 6.5. Top panel reflects the power (µW) needed to maintain isothermal conditions. Bottom panel reflects the integration of each injected peak to provide the associated energy (kJ/mol) from each injection. A line of best fit is shown. The first peak was not integrated. The reaction thermodynamics are tabulated in **Table 1**. **C)** *At*NPL4 titrated against *At*UFD1B. **D)** *At*CDC48A titrated against *At*NPL4 (left) and *At*CDC48A titrated against *At*UFD1B (right). **E)** *At*CDC48A saturated with *At*NPL4 titrated against *At*UFD1B. **F)** Unfolding of Ub-GFP by *At*CNU and *Hs*CNU complexes. *Left panel:* Unfolding was measured as a function of normalised fluorescence for *At*CNU with Ub-GFP (light blue), crosslinked *At*CNU with Ub-GFP (dark blue), and *Hs*CNU with Ub-GFP (pink) measured over time. The fluorescence was normalised to Ub-GFP alone (grey). Representative traces are shown (n = 3). *Right Panel:* Unfolding was measured as a function of normalised fluorescence for *At*CDC48–*At*UFD1B with Ub-GFP (purple), *At*CDC48–*At*NPL4 with Ub-GFP (green), and *At*CNU with Ub-GFP (blue) measured over time. The fluorescence was normalised to *At*CDC48A alone with Ub-GFP (grey). All reactions occurred in the presence of 20S proteasome (10 nM). Representative traces are shown (n = 3).

**Table 1.** Tabulated thermodynamic data from all ITC interactions. All interactions with *At*CDC48A were in the presence of AMPPNP, unless noted by subscript ADP. An asterisk indicates that N has been set to 0.16. All interactions were performed in triplicate. Values are given as mean ± standard deviation (n=3).

|  | **N** | **K<sub>D</sub>**<br>**(μM)** | **ΔG**<br>**(kJ/mol)** | **ΔH**<br>**(kJ/mol)** | **TΔS**<br>**(kJ/mol)** |
| --- | --- | --- | --- | --- | --- |
| ***AtNPL4* – *AtUFD1B*** | N/A | N/A | N/A | N/A | N/A |
| ***AtCDC48A* – *AtNPL4*** | 0.16* | 2.6 ± 0.5 | -31.8 ± 0.5 | -90.8 ± 16.4 | -59.0 ± 16.8 |
| ***AtCDC48A* – *AtNPL4B*<sub>F105A</sub>** | 0.16* | 4.6 ± 2.1 | -30.6 ± 1.1 | -46.3 ± 2.9 | 15.8 ± 3.2 |
| ***AtCDC48A* – *AtNPL4B*<sub>UBXL</sub>** | 0.9 ± 0.1 | 3.2 ± 0.3 | -31.4 ± 0.3 | -15.5 ± 0.1 | -15.8 ± 1.4 |
| ***AtCDC48A*<sub>ADP</sub> – *AtNPL4*** | 0.16* | 7.1 ± 1.8 | -29.5 ± 0.6 | -67.6 ± 13.9 | 38.1 ± 13.3 |
| ***AtCDC48A* – *AtUFD1B*** | 0.16* | 4.4 ± 1.0 | -30.6 ± 0.7 | -112.0 ± 22.3 | -81.9 ± 22.8 |
| ***AtCDC48A* + *AtNPL4* – *AtUFD1B*** | 0.16* | 3.8 ± 0.1 | -30.9 ± 0.1 | -103.6 ± 22.2 | -72.8 ± 22.2 |

**Table 2.**
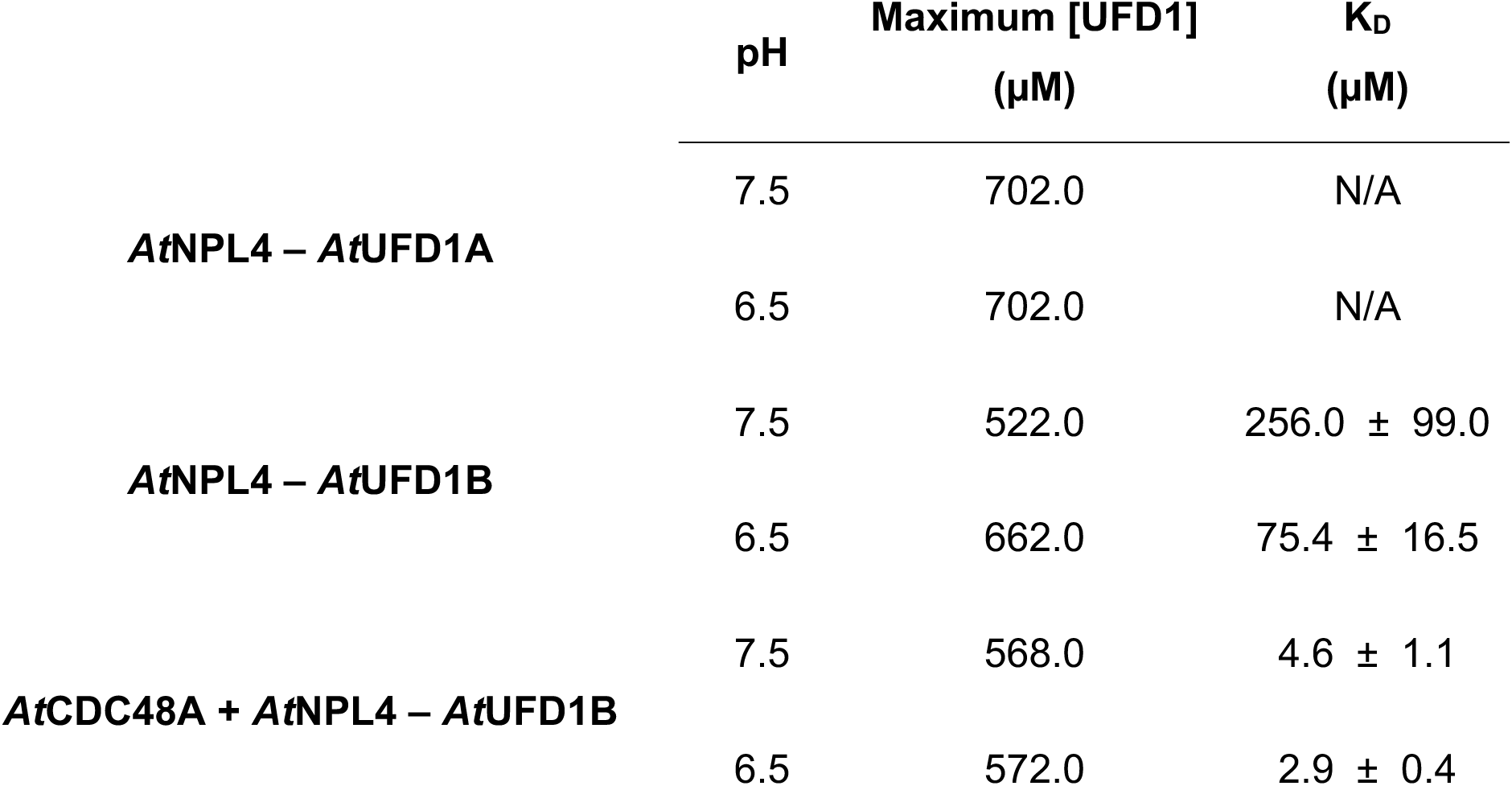
Tabulated MST interaction data. All interactions with *At*CDC48A were in the presence of AMPPNP. All interactions were performed in triplicate. Values are given as mean ± standard deviation (n=3).

| | pH | Maximum [UFD1]<br>(μM) | K<sub>D</sub><br>(μM) |
| --- | --- | --- | --- |
| **AtNPL4 – AtUFD1A** | 7.5 | 702.0 | N/A |
|  | 6.5 | 702.0 | N/A |
| **AtNPL4 – AtUFD1B** | 7.5 | 522.0 | 256.0 ± 99.0 |
| | 6.5 | 662.0 | 75.4 ± 16.5 |
| **AtCDC48A + AtNPL4 – AtUFD1B** | 7.5 | 568.0 | 4.6 ± 1.1 |
| | 6.5 | 572.0 | 2.9 ± 0.4 |

ITC analysis demonstrated that *At*CDC48A and *At*UFD1B interacted with a K_d_ of 4.4 +/- 1.0 μM, as expected by the conservation of the SHP1 motif and in agreement with previous reports on CDC48–UFD1 interactions in humans and yeast (**Figure 4D**; **Table 1**). Moreover, *At*NPL4 bound *At*CDC48A with a K_d_ of 2.6 +/- 0.5 μM, displaying an interaction stoichiometry of approximately one *At*NPL4 per *At*CDC48A hexamer (**Figure 4D**; **Table 1**). This NPL4–CDC48 interaction in absence of UFD1 contrasts with the human orthologues, where *Hs*Npl4 reportedly does not bind to *Hs*p97 without *Hs*Ufd1 (Lass et al., 2008).

To investigate whether *At*NPL4 and *At*UFD1 bind cooperatively to *At*CDC48A, we saturated *At*CDC48A with *At*NPL4 and then analysed the interaction with *At*UFD1B using MST and ITC. Our results showed no significant difference in *At*UFD1B binding to *At*CDC48A, regardless of *At*NPL4 saturation (**Figure 4E**; **Table 1**; **Table 2; Supplemental Figure 17B**). This suggests that *At*NPL4 and *At*UFD1 interacted independently with *At*CDC48A. SEC-MALS experiments in the presence of AMPPNP and size-exclusion chromatography coupled with small-angle X-ray scattering (SEC-SAXS) experiments in the absence of nucleotide further confirmed the lack of a stable cooperative complex. The preformed 6:1:1 tripartite complex of *At*CDC48A, *At*UFD1B and *At*NPL4 dissociated, with each protein eluting as a distinct peak. Additionally, no significant differences were found in the predicted molecular weight of the *At*CDC48A peaks with or without cofactors (**Figure 4A and 4B; Supplemental Figure 18**). Thus, we concluded that UFD1 and NPL4 interact independently with CDC48 in *Arabidopsis*. By carefully optimising crosslinking using 0.025% glutaraldehyde, we generated a stable complex of 620 kDa, consistent with six *At*CDC48A and one copy each of *At*UFD1B and *At*NPL4 (**Figure 4A and 4B**).

We next used an *in vitro* assay to assess the unfolding of poly-ubiquitinated GFP (Ub-GFP) by the Arabidopsis CDC48A–NPL4–UFD1B (CNU) complex (Wang et al., 2024). The assay demonstrated that the *At*CNU complex, but not *At*CDC48A alone, could unfold the ubiquitinated target *in vitro*, at a rate comparable to the human CNU complex (**Figure 4F**). Furthermore, using the mildly crosslinked complex, we established that crosslinking did not compromise its unfolding capability, validating this method for subsequent biochemical and structural investigations (**Figure 4F**).

Additionally, we compared the ATP hydrolysis rates of the plant and human CNU complexes. In the presence of a ubiquitinated substrate, the *At*CNU complex hydrolysed more ATP molecules in an endpoint assay than its human counterpart (**Figure 2D**).

Importantly, even *At*NPL4 or *At*UFD1 alone enabled *At*CDC48 to unfold ubiquitinated GFP, in line with their individual and independent association with *At*CDC48A. However, the presence of both cofactors led to a greater unfolding efficiency, inferring synergy (**Figure 4F**).

### Structure of the *At*CNU complex reveals plant-specific cofactor interactions

We next used cryo-EM to visualise the *At*CNU complex. Given that the tripartite complex dissociated during SEC, but remained functional when mildly crosslinked, we used the crosslinked sample for cryo-EM. This approach allowed us to determine the structure of *At*CNU at a resolution of 3.9 Å. The overall architecture was similar to the human and yeast counterparts, featuring a hexameric CDC48 arrangement with an additional density on one NTD and a prominent structure extending slightly off-centre above the D1 domain (**Figure 5A**).

**Figure 5.**
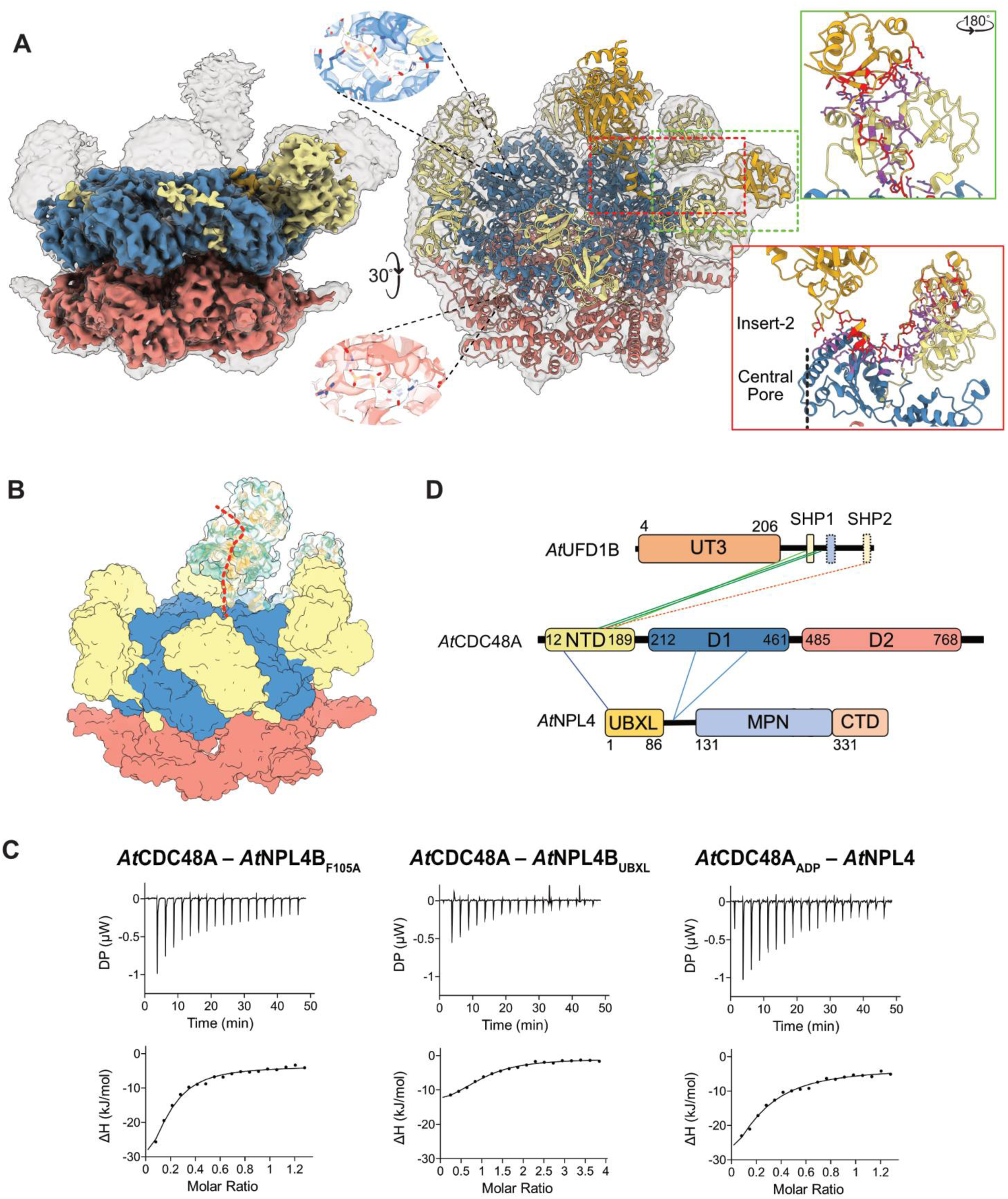
Structure of the *At*CNU complex. **A)** Cryo-EM reconstruction of *At*CNU. The unsharpened map of *At*CNU (**EMD-63612**) is overlaid as a transparent surface over its atomic model (**PDB 9M3Z**). The *At*CDC48A NTD, D1, and D2 are coloured yellow, blue, and red, respectively. *At*NPL4 is coloured orange. Spotlights show the nucleotide binding domains of D1 (top) and D2 (bottom). The red outlined box is a magnified view of the red dashed box, with the *At*CDC48A–*At*NPL4 interacting residues coloured red (*At*NPL4) and purple (*At*CDC48A). *At*NPL4’s insert-2 region and the central pore of the hexamer are labelled. The green outlined box is a magnified view of the green dashed box, with the *At*CDC48A–*At*NPL4 interacting residues coloured red (*At*NPL4) and purple (*At*CDC48A). **B)** Surface representation of *At*CNU (**PDB 9M3Z**). *At*CDC48A domains are coloured as in Panel A, and *At*NPL4 is coloured by hydrophobicity (blue–gold for hydrophilic–hydrophobic residues). The red dashed line indicates the proposed interaction site of unfolded ubiquitin along *At*NPL4 for insertion into the pore of *At*CDC48A. **C)** Representative isothermal titration calorimetry (ITC) interaction plots of AMPPNP-bound *At*CDC48A titrated against *At*NPL4B_F105A_ (left) and *At*NPL4B_UBXL_ (right), and ADP-bound *At*CDC48A_ADP_ titrated against wild-type *At*NPL4 (right). The reaction thermodynamics are tabulated in **Table 1**. **D)** Intermolecular crosslinks for *At*CDC48A– *At*UFD1B (top) and *At*CDC48A–*At*NPL4 (bottom) from crosslinking mass spectrometry. Complexes were crosslinked using the linker BS3. Lines connect the crosslinked residues. Solid lines indicate high confidence in the interaction, dashed lines indicate lower confidence in the interaction. For *At*CDC48A– *At*UFD1B, peptides including the SHP1 motif are coloured light green, peptides supporting the SHP1 interaction are coloured dark green, and peptides including the SHP2 motif are coloured orange. For *At*CDC48A–*At*NPL4, peptides including the UBXL domain are coloured dark blue and peptides including the UBXL–MPN linker are coloured light blue. Domain residues are numbered. All intermolecular crosslinked peptides for *At*CDC48A–*At*UFD1B and *At*CDC48A–*At*NPL4 are provided in **Supplemental Tables 3 and 4**, respectively.

Most of the density that was not occupied by *At*CDC48A was confidently assigned to *At*NPL4. Based on orthologous CNU complexes and AlphaFold predictions, we interpreted the density above D1 as the *At*NPL4 MPN domain, and the additional density on the nearest NTD was attributed to *At*NPL4’s UBXL domain (**Figure 5A**). Additionally, a connecting density was apparent that bridged the UBXL and MPN domains (**Figure 5A**). This region was not visible in cryo-EM maps from other organisms but matched well the UBXL–MPN linker region (residues Y87–N118) in the AlphaFold prediction of the *Arabidopsis* CDC48A and NPL4 interaction (**Figure 3C**).

The UBXL–MPN linker interacted with the hydrophobic triple-F motif of D1 (residues F268–I272), forming a strand that completed an *At*CDC48A β-sheet and a six-residue ⍺-helix (V111–R117) near the MPN domain (**Figure 5A**). This region on *At*CDC48 corresponds to the ZnF binding site in human and yeast complexes (Bodnar et al., 2018; Twomey et al., 2019; Pan et al., 2021a; Lee et al., 2023). Consistent with AlphaFold predictions, residue F105 in *At*NPL4 formed a prominent interaction with a hydrophobic pocket between the NTD and D1 (residues K67, D203, N263) (**Figure 5A**). We confirmed the UBXL–MPN linker interaction by showing that the thermodynamics of binding to AMPPNP–saturated *At*CDC48 of the F105 mutant (*At*NPL4B_F105A_) and the isolated UBXL domain (*At*NPL4B_UBXL_) were significantly different than wild-type *At*NPL4 (**Table 1**; **Figure 5C**). For these ITC experiments, we used a second variant, *At*NPL4B, that shares 84% sequence identity and conservation of F105 and all but three *At*CDC48A-interacting residues with *At*NPL4 (**Supplemental Figure 19**). Therefore, the thermodynamic parameters measured are likely to be comparable. The cryo-EM structure and structural modelling showed that the UBXL–MPN linker is too short to reach *At*CDC48A D1 in the ADP-promoted NTD-down state. In agreement, *At*NPL4 binding was reduced to 7.1 ± 1.8 µM when *At*CDC48A was saturated with ADP (*At*CDC48A_ADP_) and the coplanar NTD positioning inhibited the UBXL–MPN linker interaction (**Table 1**; **Figure 5C**). The MPN domain also engaged with D1 at the insert-2 region through a hydrophobic β-hairpin (residues V304–L311) with V308 at the tip of this turn (**Figure 5A**). Collectively, these interactions likely position the conserved *At*NPL4 groove in alignment with the central pore of the complex (**Figure 5B**).

*At*UFD1B was not resolved in the cryo-EM reconstruction due to its short interaction motifs being within an intrinsically disordered region and its ability to bind to any of the flexible NTDs of the *At*CDC48A hexamer. Weak density signals on some NTDs suggested interactions with the SHP motifs of *At*UFD1 but were insufficient for accurate modelling. However, crosslinking mass spectrometry and AlphaFold predictions supported the interaction of *At*UFD1 with the NTD of *At*CDC48A primarily through the SHP1 motif (**Figure 3D**; **Figure 5D**). Thus, the absence of unambiguous density for *At*UFD1 may be the result of averaging-out the individual interactions between the *At*UFD1 SHP1 motif and one of the six NTDs.

The *At*CNU complex was formed in the presence of AMPPNP, mildly crosslinked, then purified through SEC with ATP in the buffer (see **Methods**). Our *At*CNU structure revealed a uniform nucleotide state within D1 and D2. Each D1 active site contained either an ATP or AMPPNP molecule, consistent with the flexible upward positioning of the NTDs (**Figure 5A**). Conversely, ADP was observed in the D2 active sites, where the neighbouring arginine finger was absent and the C-terminal extension was visible on the adjacent protomer, supporting an ADP-bound configuration. Comparison with our cryo-EM structures of *At*CDC48A alone further confirmed the nucleotide state of each domain.

A 3DVA analysis identified substantial flexibility in an NTD opposite the one interacting with *At*NPL4 (**Movie 4**). Subsequent classification revealed various states where this NTD was oriented either downward or upward, with the upward orientation being more prevalent and selected for in-depth analysis.

### Features of the *At*CNU complex are conserved in plants and may be the ancestral form

We noted several characteristics unique to the *Arabidopsis* CNU: *At*NPL4 and *At*UFD1 do not form a heterodimer, interact independently with *At*CDC48A, and *At*NPL4 lacks the ZnF domains found in other eukaryotes, leading to a distinct interaction pattern with *At*CDC48A. To understand the evolutionary background of these differences, we analysed eukaryotic supergroups (**Figure 6A**). We conducted BLASTp searches using human counterparts (*Hs*Ufd1, *Hs*Npl4, and *Hs*p97) as references, focusing on species containing orthologs for all three proteins (**Figure 6B**).

**Figure 6.**
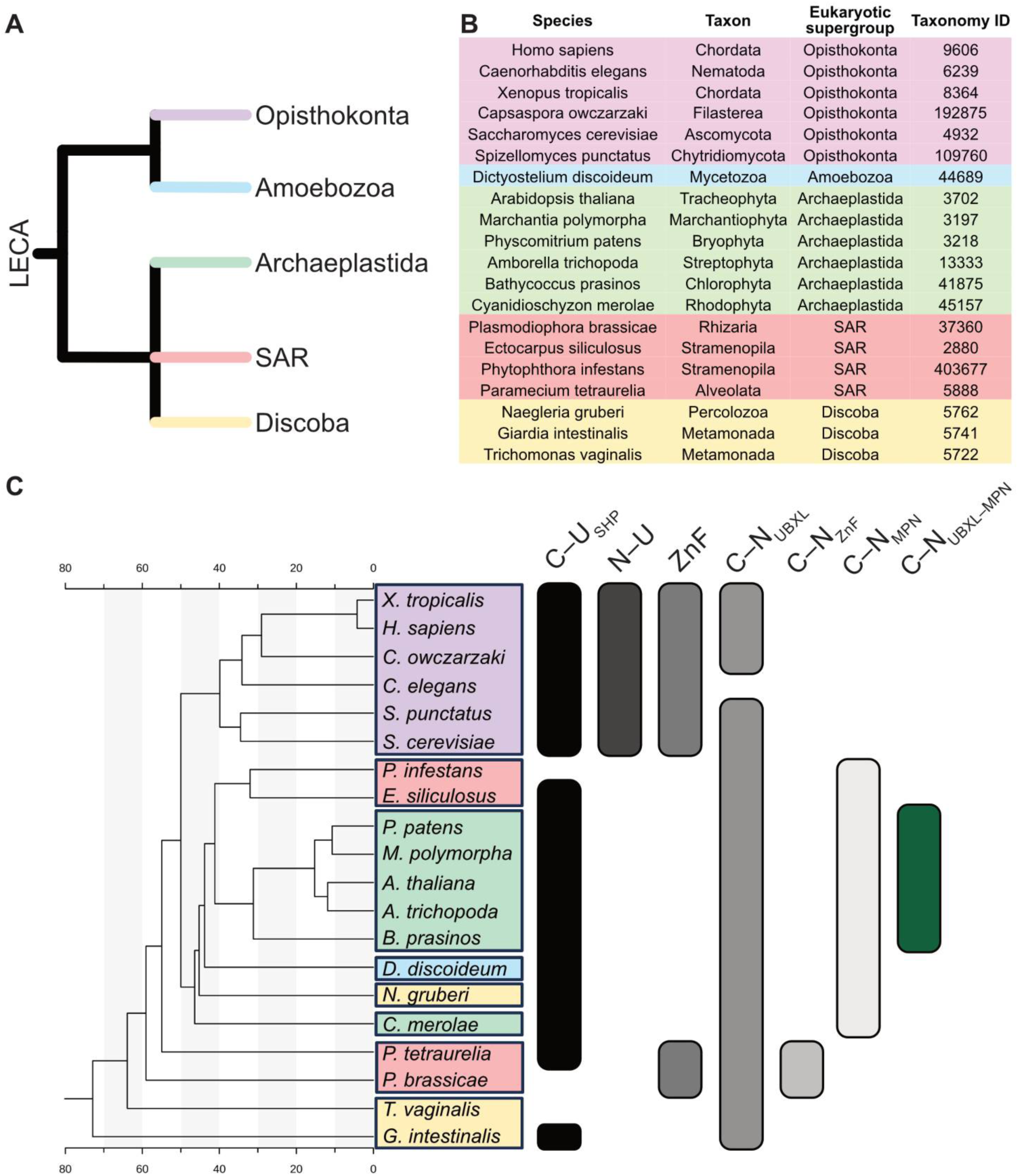
Evolutionary insights into CNU interactions. **A)** Relation of eukaryotic supergroups relative to the last eukaryotic common ancestor (LECA). Branches are coloured by eukaryotic supergroup. **B)** List of species and their taxonomic information for those used in Panel C. Adapted from (van Hooff et al., 2019), with the inclusion of additional model species and the exclusion of those with no orthologs for all of CDC48, NPL4, and UFD1. Species are coloured as in Panel A. **C)** Concatenated phylogenetic tree of CDC48, NPL4, and UFD1 with domain and interaction information from InterPro and AlphaFold multimer. Scale provided to estimate branch lengths. C–U_SHP_ refers to a predicted interaction of UFD1 to CDC48 through UFD1’s SHP motif. N–U refers to a predicted interaction through NPL4’s UBR and UFD1’s NBM. ZnF refers to NPL4 having a ZnF domain. C–N_UBXL_ refers to a predicted interaction through NPL4’s UBXL domain. C–N_ZnF_ refers to a predicted interaction through NPL4’s ZnF domain. C–N_MPN_ refers to an interaction through NPL4’s MPN domains. C-N_UBXL–MPN_ refers to a predicted interaction through NPL4’s MPN–UBXL linker region. Sequences were aligned and concatenated using MEGA11and the phylogenetic tree calculated using the UPGMA algorithm with 1000 bootstrap replicates in TreeViewer. Species are coloured as in Panel A.

We used Uniprot and AlphaFold to obtain domain and functional information from species across the eukaryotic supergroups (**Supplemental Figures 13, 14 and 16**). Using sequence alignments for each protein, we constructed a concatenated phylogenetic tree to infer the evolutionary history of CNU complex interactions (**Figure 6C**).

Our analysis showed the CDC48–UFD1 interaction was conserved in all species with UFD1 SHP motifs (**Supplemental Figure S14**). In higher eukaryotes, particularly within the Opisthokonta supergroup, NPL4 contained a ZnF domain. Additionally, these eukaryotes developed a direct UFD1–NPL4 interaction (**Supplemental Figure 15**) and a potential dependence on this heterodimer for interaction with CDC48, as indicated by an absence of ZnF interaction (**Supplemental Figure 16**). Further, our findings indicated that although the NTD–UBXL interaction between CDC48–NPL4 is conserved across nearly all studied eukaryotes (*Caenorhabditis elegans’* NPL4 lacks the UBXL domain), those without NPL4 ZnF domains utilise a short β-strand and ⍺-helix motif to anchor the MPN domain to D1 (**Supplemental Figure 16)**. In Viridiplantae in particular, the MPN anchoring was further strengthened by incorporating the UBXL–MPN linker interaction (**Figure 6C**). Taken together, our analysis suggested that the independent interactions of CDC48 with UFD1 and NPL4 may reflect an ancestral state.

## DISCUSSION

Plant-specific adaptations of the CNU degradation machinery are currently poorly understood. By combining structural, functional and computational methods, we established the molecular architecture of the *At*CNU system. As expected from the high sequence identity of CDC48 across eukaryotes, the hexameric structure of *At*CDC48A closely resembles fungal and animal orthologues. Nonetheless, we observed plant-specific features for this protein. These include differences in the dynamics of *At*CDC48A, such as the NTD position and the absence of a D2 rotation during the catalytic cycle. Our variability analysis of AMPPNP-bound *At*CDC48A suggests the removal of N-terminal extension from the neighbouring D1 and movement of the NTD allows the neighbouring D1 to reposition and sequentially coordinates the NTD position on adjacent protomers. Additionally, our structural analysis suggested that F363, part of the short region of homology involved in inter-subunit signalling, does not participate in rotamer switching following hydrolysis of ATP to ADP, as observed in orthologues <u>(Shein et al., 2024)</u>^28^. This lack of rotamer switching is a phenotype seen in Inclusion body myopathy associated with Paget disease of the bone and frontotemporal dementia (IBMPFD) mutants of *Hs*p97, which causes a dysregulation in NTD positioning such as the NTDs being locked in a Down position in the presence of an ATP analogue, or Up in the presence of ADP (Tang and Xia, 2013; Shein et al., 2024). In both the X-ray and cryo-EM structures of *At*CDC48A, we did not see the same correlation between the rotamer state of F363 and the NTD positioning (**Supplemental Figure 20**). However, our structural analysis cannot exclude transient rotamer switching during hydrolysis, and investigation using nuclear magnetic resonance or molecular dynamics simulations may provide further insights (Rydzek et al., 2020; Shein et al., 2024).

Moreover, neither the X-ray nor the cryo-EM structures of *At*CDC48A contained the same D2-meditated dodecameric architectures that were reported for *Hs*p97. The *Hs*p97 hexamer–hexamer interface, strongly involving the D2 ⍺9 helix is well conserved in *At*CDC48A (79% sequence identity for interface residues), with only the conserved E765 in *Hs*p97 replaced with Q760 in *At*CDC48A. Therefore, we cannot rule out that *At*CDC48A dodecamers were disrupted or discarded by sample preparation protocol (see **Methods**). Conversely, *At*CDC48A formed NTD–D1-mediated dodecamers in the crystallographic lattice (**Supplemental Figure 12B**), which we did not observe in cryo-EM, suggesting these dodecamers were stabilised by the crystallisation conditions.

More striking differences, however, were found in the NPL4 and UFD1 cofactors and their linkage to CDC48 in plants. Unlike in animals and fungi, *At*NPL4 and *At*UFD1 do not associate with each other to bind CDC48 jointly. Instead, each cofactor binds independently to *At*CDC48A. The binding of cofactors positioned most of the NTDs in an Up conformation, except for the opposite NTD which remained flexible. Each cofactor interacted with a stoichiometry of one cofactor to one hexamer of *At*CDC48A. This stoichiometry likely results from steric hinderance over the D1 domain preventing additional copies from attaching, as we saw a stoichiometry of six *At*NPL4B_UBXL_ per hexamer when the obstructing MPN domain and UBXL–MPN linker were absent. The highly disordered *At*UFD1 likely obstructs other copies in a similar manner, however, its flexibility may allow for interacting with an already formed *At*CDC48A–*At*NPL4 complex.

The individual interactions are only partly similar to those in animals and fungi in that the AtCDC48 NTDs bind to the conserved SHP motifs of AtUFD1 and the UBXL domain of AtNPL4. However, plant NPL4 lack the ZnF domains that NPL4 of other eukaryotic lineages use to bind to CDC48 D1. *At*NPL4 compensates for this lack through a plant-specific UBXL–MPN linker that flexibly tethers the MPN domain near the *At*CDC48 pore, in a position akin to its animal and yeast orthologues. However, the MPN domain of AtNPL4 has a shortened insert-2 region, which in animals and fungi protrudes over the pore in substrate-free complexes and away from the pore in substrate-bound complexes (Twomey et al., 2019; Pan et al., 2021a; Lee et al., 2023). Thus, it is possible that plants employ a different substrate initiation mechanism compared to orthologs.

The lack of direct interaction between *At*NPL4 and *At*UFD1 indicates they do not form an obligate heterodimer in plants, conversely to animals and fungi (Sato et al., 2019; Nguyen et al., 2022). Indeed, although each interaction was not very strong, either cofactor alone was sufficient to induce unfolding through *At*CDC48A alone, but at a greater rate when combined. The distinct *At*CDC48A domain dynamics and the absence of ZnF domains in *At*NPL4 and the alternative binding mechanisms may also reflect this cofactor independence in plants. Our bioinformatic analyses suggest that the independent binding of *At*NPL4 and *At*UFD1 represent an ancestral state, with the obligate heterodimer seen in animal and yeast being a derived characteristic. Although the absence of ZnF domains in *At*NPL4 would limit its regulation through metal ions (Skrott et al., 2017; Pan et al., 2021a), the independent and modular association of NPL4 and UFD1 may present advantages for plants. For example, by enabling independent regulatory mechanisms for NPL4 and UFD1, and by allowing a greater combinatorial space of cofactors for *At*CDC48A, the increased modularity may provide a more flexible and adaptable substrate processing system in plants, especially if cofactors of the plant UBX (PUX) family could also be combined with NPL4 (Zhang et al., 2022; Inès et al., 2024). This modularity may reflect the cellular demands and environmental challenges of sessile plants, and hence be particularly important for processes such as germination, development, flowering, regulation of chloroplast function, and plant immunity.

## METHODS

### Protein Expression and Purification

The cDNA sequences for *At*CDC48A (AT3g09840), *At*NPL4 (AT2g47970), *At*UFD1A (AT2g29070), and *At*UFD1B (AT2g21270) were purchased from Integrated DNA Technologies (IDT) and cloned into the pGEX6P-1 vector between the BamHI and NotI restriction sites. Variants *At*CDC48A_NTD_ (Residues 28-190) and *At*CDC48A_28-809_ were cloned into the pGEX6P-1 vector between the BamHI and NotI restriction sites using restriction primers. *At*NPL4B_UBXL_ (AT3g63000; Residues 1-89), *At*NPL4B_F105A_ ( AT3g63000), *Hs*p97 (HGNC:12666), *Hs*Npl4 (HGNC:18261), and *Hs*Ufd1 (HGNC:12520) were ordered from Twist Biosciences in the pGEX6P-1 vector. All proteins were expressed in *Escherichia coli* BL21 (DE3) cells with an N-terminal GST tag and a PreScission protease cleavage site. Proteins were expressed and purified following the optimised protocol in (Huntington et al., 2022). Briefly, cells were cultured in the presence of ampicillin, expression was induced using isopropyl β-D-1-thiogalactopyranoside (IPTG), and pellets were stored at −80°C for later purification. For purification, pellets were resuspended in lysis buffer (50 mM Tris pH 7.5, 200 mM NaCl, 3 mM dithiothreitol (DTT), one tablet of complete protease inhibitor (Roche) per 50 mL preparation, and 0.5% Triton X-100), incubated with benzonase nuclease (Millipore) and lysed by sonication. The lysate was clarified through centrifugation and purified by application to a GST-affinity column twice, with high salt washes after each application. The proteins were cleaved and further purified through SEC. An additional purification step was conducted for *At*CDC48A_NTD_, where the peak fraction from SEC was loaded onto a MonoQ column with a low salt (50 mM NaCl) SEC buffer and eluted along a NaCl concentration gradient. Protein purity was assessed by SDS-PAGE, and relevant fractions were pooled and stored at −80°C or used fresh, as described.

### Cryo-Electron Microscopy

#### Sample preparation

For the *At*CDC48A nucleotide states, nucleotide-free *At*CDC48A was freshly purified (Huntington et al., 2022), the peak hexameric fraction retained without concentrating, 2 mM of adenosine diphosphate (ADP) or adenylyl imidodiphosphate (AMPPNP) was added with 2 mM MgCl2, and the mixture incubated on ice for 30 minutes. No nucleotide or MgCl_2_ was added for apo *At*CDC48A.

The *At*CNU complex was prepared at an equimolar protein concentration with 2 mM AMPPNP and 2 mM MgCl_2_ and incubated on ice for 30 minutes. The complex was crosslinked with 0.025% glutaraldehyde for 10 minutes at room temperature and the reaction was quenched by adding ⅕ volume of 1 M glycine. The complex was purified on an analytical 10/300 Superdex 200 column (GE Healthcare) using SEC buffer with an additional 100 µM ATP and 2 mM MgCl_2_. The peak fraction was retained without concentrating and stored at −80°C.

#### Specimen preparation

Cryo-EM specimens were prepared using the Vitrobot Mark IV (Thermo Fisher Scientific) at 22°C with 100% humidity. Samples were prepared by applying 2-3 μL of protein solution and blotting for 3 s. Samples of apo *At*CDC48A (2 mg/mL) were prepared on glow-discharged Cu grids (C-flat Cu 300 mesh R1.2/1.3, Protoships Inc.). Samples of AMPPNP bound *At*CDC48A (2 mg/mL), ADP-bound *At*CDC48A (2 mg/mL), and the *At*CNU complex (1.5 mg/mL) were prepared on glow-discharged Au grids (C-flat Au 300 mesh R1.2/1.3, Protoships Inc.).

#### Data collection

Cryo-EM data were collected at multiple facilities. At KAUST, datasets for apo and AMPPNP-bound *At*CDC48A were recorded on a Titan Krios G2 (ThermoFisher Scientific) operated at 300 kV, equipped with a K2 Summit direct detector and a GIF BioQuantum imaging filter (Gatan). Movies were collected at 130k magnification in super-resolution mode, yielding a pixel size of 0.52 Å/pixel at the specimen level. For the apo *At*CDC48A dataset, 1,097 MRC movies were collected with a total fluence of 50 e^-^/Å^2^, using Latitude S with a defocus range of −2.2 µm to −1.0 µm in 0.3 µm intervals. For the AMPPNP-bound AtCDC48A dataset, two grids were collected under similar conditions: 1,095 MRC movies from Grid 1 and 2,284 MRC movies from Grid 2. At scopeM, ETH Zürich, datasets for ADP-bound *At*CDC48A and the *At*CNU complex were recorded on a Titan Krios G3 (ThermoFisher Scientific) equipped with a K3 direct detector and GIF BioQuantum imaging filter (Gatan). For the ADP-bound *At*CDC48A dataset, 16,380 TIFF movies were collected at 130k magnification, yielding a pixel size of 0.66 Å/pixel at the specimen level and a total fluence of 64 e^-^/Å^2^, using aberration-free image shift (AFIS) in EPU with a defocus range of −2.6 µm to −0.8 µm in 0.2 µm intervals. For the AtCNU dataset, 15,984 TIFF movies were collected under identical conditions, with a total fluence of 66 e^-^/Å^2^.

#### Data processing

A detailed data processing scheme for each dataset can be found in the supplemental materials (**Supplemental Figures 2, 4, 7 and 9**). The general processing scheme consisted of the following steps. Movies were motion-corrected in RELION (Kimanius et al., 2021) and transferred to CryoSPARC (Punjani et al., 2017) for picking using Topaz (Bepler et al., 2020) or Gautomatch (Gautomatch - SBGrid Consortium - Supported Software) and iterative 2D classification for particle curation. Ab initio Reconstruction was then performed to generate junk decoy classes and high-resolution references. These were used together with an over-picked particle stack for particle curation in 3D using iterative Heterogeneous Refinement jobs in which only particles retained in the target classes were used in the next iteration. Then, Non-Uniform Refinement (Punjani et al., 2020) was used to generate a consensus reconstruction that was subjected to 3DVA in CryoSPARC (Punjani and Fleet, 2021) and transferred back to RELION using Pyem (Asarnow et al., 2019). The particle stacks were iteratively refined using Bayesian Polishing and CTF Refinement. Class3D without alignments was then used to generate more homogeneous particle stacks from which the final reconstructions were produced. In RELION, Blush (Kimanius et al., 2024) was used when possible. Validation metrics for the final maps can be found in the Supplemental Material (**Supplemental Figures 3, 5, 6, 8 and 10**).

#### Variability analysis

3DVA (Punjani and Fleet, 2021) was performed using a filter resolution of 6 Å and a global mask. The heterogeneity was represented along 3 eigenvectors. 3DVA Display in Simple mode was used to generate 20 volumes along each eigenvector, and movies were generated for each volume series using ChimeraX (Pettersen et al., 2021). The heterogeneity along the first and second eigenvector of AMPPNP *At*CDC48A was discretised using 3DVA Display in Cluster mode to generate 5 clusters. The clusters were then refined using Local Refinement using the same mask.

#### Model building and refinement

Starting models were generated using AlphaFold. To improve the interpretability of maps for the initial stages of model building, unsharpened maps were subjected to the EMReady algorithm (He et al., 2023). Model building and refinement of the *At*CDC48A nucleotide states were performed on one protomer and symmetrically applied to the others. Briefly, the refinement of each model consisted of the following steps. Distance restraints were imposed on the AlphaFold model to retain the local geometry of the predicted model, followed by real-space refinement inside of Coot (Emsley and Cowtan, 2004) to enable large movements and fit the domains and secondary structure elements to the coulomb potential. The nucleotides and Mg^2+^ ions were added, locally refined and manually checked for proper fit and geometry. The models were then iteratively refined using Phenix real space refine (Liebschner et al., 2019) and manual building in Coot using the unfiltered primary map deposited.

### AlphaFold Structure Prediction

AlphaFold predicted structures and modelling of the interactions was performed using an in-house wrapper (Guzmán-Vega and Arold, 2024) for AlphaFold2 multimer (Jumper et al., 2021; Evans et al., 2022).

### Crosslinking Mass Spectrometry

*At*CDC48A–AtNPL4 and *At*CDC48A–*At*UFD1B were crosslinked by incubating *At*CDC48A (5 µM) with *At*NPL4 or *At*UFD1B (10 µM), followed by addition of 0.1 mM bissulfosuccinimidyl suberate (BS3) for 20 minutes at 25°C. The reaction was quenched with 100 mM Tris-HCl (pH 7.5). Crosslinked proteins were separated on a 4–12% SDS-PAGE gel, and target bands were excised, cut into 1 mm pieces, and washed sequentially with 100 mM and 25 mM ammonium bicarbonate/acetonitrile (1:1) for destaining. Gel pieces were dehydrated with acetonitrile, reduced with 10 mM DTT (56°C, 60 minutes), and alkylated with 55 mM iodoacetamide (room temperature, 30 minutes). After washing, proteins were digested with trypsin (1 µg/µL in 25 mM ammonium bicarbonate, 60 minutes on ice). Peptides were extracted three times with 50% acetonitrile/0.1% formic acid, concentrated, and resuspended in 20 µL of 0.1% formic acid for LC-MS.

Liquid-chromatography mass spectrometry (LC-MS) analysis was performed using an Orbitrap Fusion Lumos mass spectrometer coupled to a Dionex UltiMate 3000 RSLC nano HPLC system. Peptides were loaded onto an Acclaim PepMap 100 C18 trap column (100 µm × 2 cm, 100 Å) and separated on a PepMap RSLC C18 analytical column (2 µm, 100 Å, 75 µm × 50 cm) at 300 nL/min over a 75-minute gradient. Mobile phases were 0.1% formic acid in water (solvent A) and 0.1% formic acid in 95% acetonitrile (solvent B). Peptides were eluted using a 55-minute gradient (2.1% to 31.6% B), followed by a 5-minute increase to 90% B, held for 3 minutes, and re-equilibrated at 2.1% B for 10 minutes. Ionization was performed at 2 kV using an EASY-Spray emitter. Data were acquired in data-dependent mode with a 3-second cycle time. Survey scans (m/z 400–1600) were performed at 120,000 resolution, and the most intense ions (charge states 2+ to 6+) were fragmented via HCD (30% collision energy, 1.6 Th isolation window). Fragment spectra were acquired at 15,000 resolution.

Peak lists in mgf format were searched using Protein Prospector ‘Batch-Tag Web’ (Trnka et al., 2014) with a charge range of 2-5, a precursor tolerance of 20 ppm, and a fragment ion tolerance of 1 Da. The instrument setting was defined as ESI-Q-hi-res. Trypsin was specified as the cleavage enzyme (up to two missed cleavages), with carbamidomethylation of cysteine as a fixed modification and methionine oxidation as a variable modification. Crosslinked peptides were analysed using the ‘Search Compare’ program.

### X-ray Diffraction

Crystallisation was performed using the sitting drop vapour diffusion method with the concentrated protein at 20 mg/mL for both *At*CDC48A_28-809_ and the NTD alone. Crystals were obtained under crystallization conditions of 100mM HEPES, 20mM magnesium chloride hexahydrate, 22% poly acrylic acid 5100, 2mM ADP pH 7.5 for *At*CDC48A_28-809_ and 800mM Sodium phosphate monobasic, 1200mM Potassium phosphate dibasic, 100mM Sodium acetate/Acetic acid pH 4.5 for the NTD. Crystals were flash cooled in liquid nitrogen using 25% glycerol as a cryoprotectant. X-ray diffraction data were collected at the Proxima 2A beamline of the SOLEIL synchrotron. The ADP-bound *At*CDC48A_28-809_ and NTD crystals initially diffracted to resolutions of 3.4 Å and 2.6 Å, respectively. Data integration and scaling were performed using XDS. The structure was determined using the molecular replacement method implemented in PHASER (McCoy et al., 2007), with AlphaFold-generated models of *At*CDC48A_28-809_ and the NTD as search models. Iterative cycles of model building and refinement were performed using Coot (Emsley and Cowtan, 2004) and REFMAC5 (Murshudov et al., 2011) until the R and R_free_ values converged to 0.34 and 0.38 for the ADP-bound *At*CDC48A_28-809_ and 0.24 and 0.31 for the NTD. Diffraction data and refinement statistics are summarised in **Supplemental Table 2**.

### Differential Scanning Fluorimetry

Thermal unfolding profiles determined the thermal stability of *At*CDC48A in the presence and absence of different nucleotides. All assays were carried out at a final protein concentration of 5 μM, in buffer containing 50 mM Tris–HCl pH 7.5, 150 mM NaCl, 1mM TCEP, and with Sypro Orange dye (Invitrogen) at a 2x final concentration. Thermal denaturation temperatures were obtained in a CFX96 touch real-time thermal cycler (Bio-Rad) with an optical reaction module by ramping the temperature between 20 and 95°C at 1°C per minute, and fluorescence acquisition through the channel. Data were analysed with CFX Manager Software V3.0 (Bio-Rad) and plotted using PRISM (v10.0).

### Size Exclusion Chromatography–Multi-Angle Light Scattering (SEC-MALS)

All SEC-MALS experiments were conducted using an analytical Superdex S200 Increase 10/300 GL column (Cytiva) in SEC-MALS buffer (20 mM HEPES pH 7.5, 150 mM NaCl, and 3 mM DTT). Proteins and complexes were incubated on ice for 30 minutes prior to analysis. For complexes of *At*NPL4 with *At*UFD1, equimolar concentrations of each protein were used. For complexes containing *At*CDC48A, a molar ratio of 6:1:1 (*At*CDC48A:*At*NPL4:*At*UFD1B) was used, and the complex was concentrated using an Amicon-0.5 filtration device with a 30 kDa molecular weight cutoff (MWCO) before SEC-MALS analysis. Additionally, all *At*CDC48A-containing complexes were prepared with 2 mM AMPPNP and 2 mM MgCl_2_, and the running buffer was supplemented with 100 µM ATP. *At*CNU complex was prepared at a molar ratio of 6:1:1 (*At*CDC48A:*At*NPL4:*At*UFD1B) and crosslinked for 10 minutes at room temperature using a glutaraldehyde concentration of 0.05% before quenching with ⅕ volume of 1 M glycine and being applied to the SEC-MALS system. The Astra Multiangle Light Scattering software (Wyatt Technology) was used to calculate the refractive index and estimate the molecular weight.

### Microscale Thermophoresis (MST)

For all MST experiments, *At*NPL4 was labelled with Alexa-488 dye (ThermoFisher) at a final concentration 1.5 times the protein concentration. The protein-dye solution was incubated in an opaque tube while rotating for two hours at room temperature. Excess dye was removed using an Amicon-0.5 filtration device with a 10 kDa molecular weight cutoff (MWCO) by repeatedly diluting the solution with buffer until the flowthrough was free of dye. Before experiments, all proteins were dialysed into either pH 6.5 buffer (50 mM Bis-Tris pH 6.5, 100 mM NaCl, 1 mM TCEP) or pH 7.5 buffer (50 mM Tris–HCl pH 7.5, 100 mM NaCl, 1 mM TCEP), depending on the experimental design. The interacting partner (*At*UFD1A or *At*UFD1B) was serially diluted into sixteen aliquots.

For interactions between *At*NPL4 and *At*UFD1A or *At*UFD1B, the labelled protein was diluted to a dye concentration of 50 nM, and 1% Tween-20 was added. A 10 µL aliquot of this solution was mixed with each 10 µL dilution of the interacting partner. For interactions involving *At*CDC48A saturated with *At*NPL4 and *At*UFD1B, the labelled protein was diluted to a final dye concentration of 100 nM with 1% Tween-20 and 40 µM *At*CDC48A. A 5 µL aliquot of each solution was added to each 15 µL dilution series to achieve final concentrations of 25 nM dye and 10 µM *At*CDC48A. Solutions in *At*CDC48A experiments were incubated with 2 mM AMPPNP and 2 mM MgCl_2_. All interactions were measured using the Nanotemper Monolith NT.115 MST instrument with an excitation power of 25%. Triplicate measurements were taken at MST powers of 20%, 40%, and 60%, except for *At*CDC48A saturated with *At*NPL4 and *At*UFD1B at pH 7.5, in which only 20% and 40% MST powers were recorded.

### Isothermal Titration Calorimetry (ITC)

Proteins were dialysed overnight and degassed in ITC buffer (50 mM Bis-Tris pH 6.5, 100 mM NaCl, 1 mM TCEP, and 2 mM MgCl_2_). AMPPNP (Sigma) or ADP (Sigma) were dissolved in the same buffer and added at a final concentration of 2 mM to both the cell and syringe solutions and incubated at room temperature for 20 minutes. All interactions with *At*CDC48A were performed in the presence of AMPPNP unless indicated. Before each experiment, the cell and syringe were rinsed with milli-Q H_2_O and ITC buffer, and the reference cell was filled with milli-Q H_2_O.

For the interaction between *At*NPL4 and *At*UFD1B, *At*NPL4 was loaded into the cell at 10 µM, and *At*UFD1B was loaded into the syringe at 200 µM. For interactions involving *At*CDC48A (or *At*CDC48A_ADP_) with *At*NPL4, *At*NPL4B_F105A_, or *At*UFD1B, *At*CDC48A was loaded into the cell at 30 µM, and the interacting protein was loaded into the syringe at 200 µM. For the interaction between *At*CDC48A and *At*NPL4B_UBXL_, *At*CDC48A was loaded into the cell at 10 µM, and *At*NPL4B_UBXL_ was loaded into the syringe at 200 µM. For the ternary interaction involving *At*CDC48A saturated with *At*NPL4, and *At*UFD1B, the cell contained 30 µM *At*CDC48A and 60 µM *At*NPL4, while the syringe contained 200 µM *At*UFD1B. All titrations were performed at 25°C with an initial injection of 0.4 µL, followed by 18 injections of 2 µL each, using an ITC200 calorimeter (Malvern). Data analysis and figure preparation were performed using the MicroCal PEAQ-ITC Analysis Software (Malvern).

### Small-Angle X-ray Scattering (SEC-SAXS)

SAXS data were recorded at the SWING beamline (SOLEIL, Saint-Aubin, France) using a wavelength of 1.03 Å. The distance of the sample to the detector was 1.8 m, resulting in the momentum transfer of a range of 0.1 1/Å < q < 0.5 1/Å. Data acquisition was carried out by size exclusion chromatography using a Bio SEC-3 HPLC (Agilent) column coupled with SAXS at 15_°_C. Buffer data were calculated from the buffer (20 mM HEPES pH 7.5, 150 mM NaCl, 1 mM TCEP) eluted before the proteins, and subtracted from the protein data using CHROMIXS (Panjkovich and Svergun, 2018), as part of the ATSAS software package (Manalastas-Cantos et al., 2021). Samples were prepared at an equimolar concentration of 33 µM at a volume of 50 µL and incubated on ice for 30 minutes prior to injection. Complexes were formed without adding nucleotide. Estimation of the molecular weight distribution was performed using MW-DARA (Kikhney et al., 2016).

### Unfolding Assay

Unfolding assays were performed at 37°C. The reaction mixture contained 700 nM Ub-GFP substrate, 500 nM *At*CDC48A or *Hs*p97, 4 mM *At*NPL4 or *Hs*Npl4 and *At*UFD1B or *Hs*Ufd1, and either in the presence or absence of 10 nM 20S proteasome (Enzo) in unfolding assay buffer (20 mM HEPES pH 7.5, 150 mM NaCl, 10 mM MgCl₂, 5 mM ATP). Slight variations in protein concentrations were applied between assays. Proteins were mixed and transferred into a black 384-well plate with a clear bottom. Fluorescence intensity was measured from the bottom of the plate at excitation and emission wavelengths of 485 nm and 510 nm, respectively, using a TECAN Infinite® M1000 PRO microplate reader or a PHERAstar® FS microplate reader. Measurements were taken at 20-second intervals. Relative fluorescence was calculated by normalising the fluorescence signal to its initial value at time 0.

### ATPase Activity Assay

ATP hydrolysis activity was measured using the Malachite Green Phosphate Assay Kit (Sigma-Aldrich) to quantify free phosphate released from ATP hydrolysis by *At*CDC48A or p97. The assay was conducted at 30°C in ATPase buffer (50 mM HEPES (pH 7.5), 50 mM NaCl, 10 mM MgCl₂, 0.5 mM TCEP, and 0.1 mg/mL BSA). Prior to ATP addition, *At*CDC48A or *Hs*p97 (150 nM) was pre-incubated for 10 minutes with *At*NPL4 or *Hs*Npl4 (500 nM), *At*UFD1B or *Hs*Ufd1 (500 nM), and Ub-GFP (2 μM). The reaction was initiated by the addition of ATP and allowed to proceed for 30 minutes. The reaction was terminated by adding the Malachite Green reagent, followed by a 20-minute incubation. Absorbance at 640 nm was measured using a TECAN Infinite® M1000 PRO microplate reader. Phosphate concentrations were determined using a standard curve, and ATPase hydrolysis rates were calculated as the average of two or three replicates. Data were normalised relative to the hydrolysis rate of *At*CDC48A alone.

### Phylogenetic Analysis

A list of representative species from eukaryotic supergroups was adapted from (van Hooff et al., 2019) and supplemented with additional model species. The *H. sapiens* orthologues, *Hs*p97, *Hs*Ufd1, and *Hs*Npl4, were used as query sequences for BLASTp (Altschul et al., 1990) search of non-redundant protein sequences from these representative eukaryotic species. Species that did not contain orthologues for all three proteins were excluded from the analysis. Multiple sequence alignment (MSA) of each protein was performed using ClustalW, and the MSAs for each protein concatenated within MEGA11 (Tamura et al., 2021). A concatenated phylogenetic tree was then calculated using the unweighted pair-group method with arithmetic means (UPGMA) with 10000 bootstrap replicates and visualised using Treeviewer (Bianchini and Sánchez-Baracaldo, 2024). Monomeric structures of each protein and pairwise CNU complex interactions were predicted using AlphaFold. Interpro, Uniprot, MSA, and the predicted structures were used to curate a list of features for the proteins and their predicted interactions, and the concatenated tree was used to infer evolutionary relationships. The list of species and sequences used is provided in **Supplemental Table 5.**

## DATA AVAILABILITY

Cryo-EM reconstructions and atomic models have been deposited in the Electron Microscopy Data Bank (EMDB) and Protein Data Bank (PDB), respectively, under the following accession numbers: **EMD-63608**, **PDB 9M3V** (apo); **EMD-63609**, **PDB 9M3W** (AMPPNP-Up); **EMD-63610**, **PDB 9M3X** (AMPPNP-Down); **EMD-63611**, **PDB 9M3Y** (ADP); **EMD-63612**, **PDB 9M3Z** (*At*CNU). Crystallographic atomic models have been deposited in the PDB under the following accession numbers: **PDB 9M4G** (*At*CDC48A_NTD_) and **PDB 9M4N** (*At*CDC48A_28-809_)

## AUTHOR CONTRIBUTIONS

Conceptualization: **BH**, **UFSH**, and **STA**; Formal analysis: **BH**, **AS**, **JW**; Investigation: **BH**, **AS**, **JW, JZ**; Resources: **BH**, **JZ**, **LZ**, **BMQ**; Writing—original draft: **BH** and **STA**; Writing—review and editing: **BH** and **STA**; Visualization: **BH**; Supervision: **UFSH** and **STA**; Project administration: **STA**; Funding acquisition: **STA**. All authors read and approved the manuscript.

## ACKNOWLEDGMENTS

We would like to thank the Imaging and Characterization core lab, the Supercomputing Laboratories, and Bioscience core labs at KAUST for the use of their resources and support. We are grateful to ETH Zürich for access to the computational resources at ScopeM and Euler cluster.

## DECLARATION OF INTERESTS

The authors declare that the research was conducted in the absence of any commercial or financial relationships that could be construed as a potential conflict of interest.

During the preparation of this work the authors used Copilot to improve the language in some sections. After using this tool/service, the authors reviewed and edited the content as needed and take full responsibility for the content of the publication.

## FIGURE LEGENDS

**Movie 1: 3DVA of ANPPNP-bound AtCDC48A along the first eigenvector**

**Movie 2: 3DVA of ANPPNP-bound *At*CDC48A along the second eigenvector**

**Movie 3: 3DVA of apo *At*CDC48A**

**Movie 4: 3DVA of *At*CNU**

## SUPPLEMENTAL DATA

**Figure S1.**
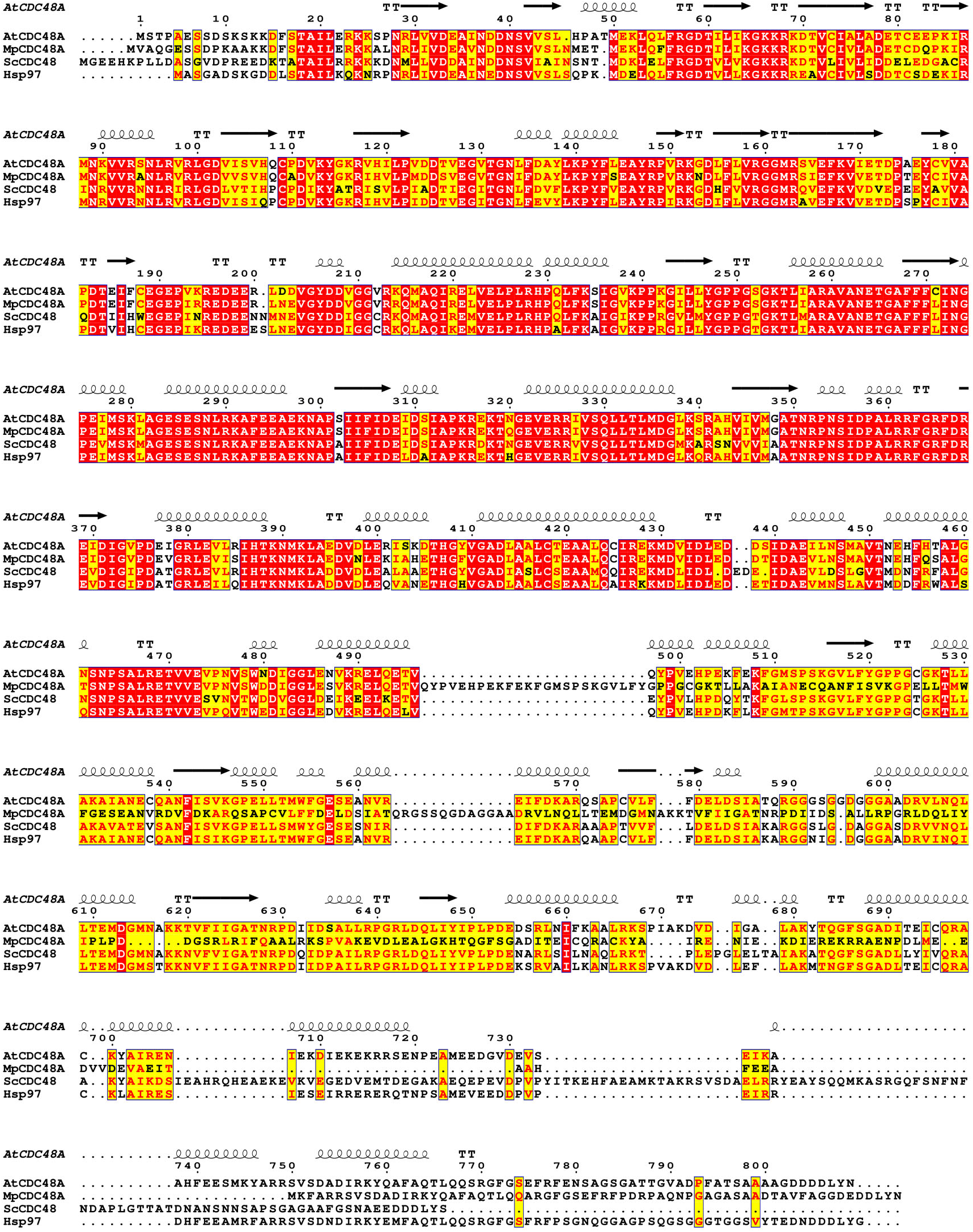
Sequence alignment of CDC48 in *Arabidopsis thaliana* (*At*), *Marchantia polymorpha* (*Mp*), *Saccharomyces cerevisiae* (*Sc*) and *Homo sapiens* (*Hs*). Secondary structure features and residue numbers are provided for *At*CDC48A above the alignment. Red colouring indicates identical residues, and yellow colouring indicates highly conserved residues.

**Figure S2:**
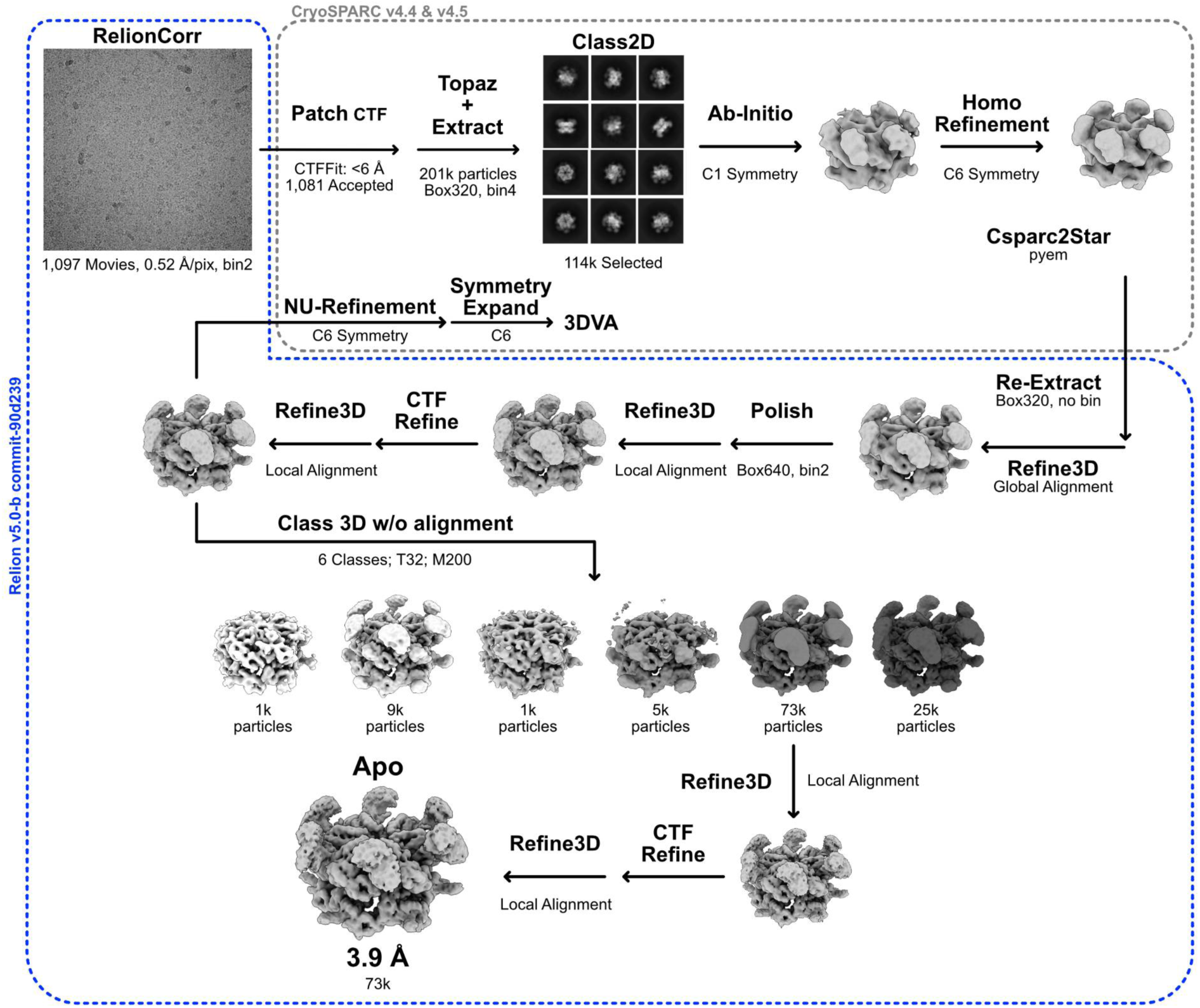
Cryo-EM image processing of apo *At*CDC48A.

**Figure S3:**
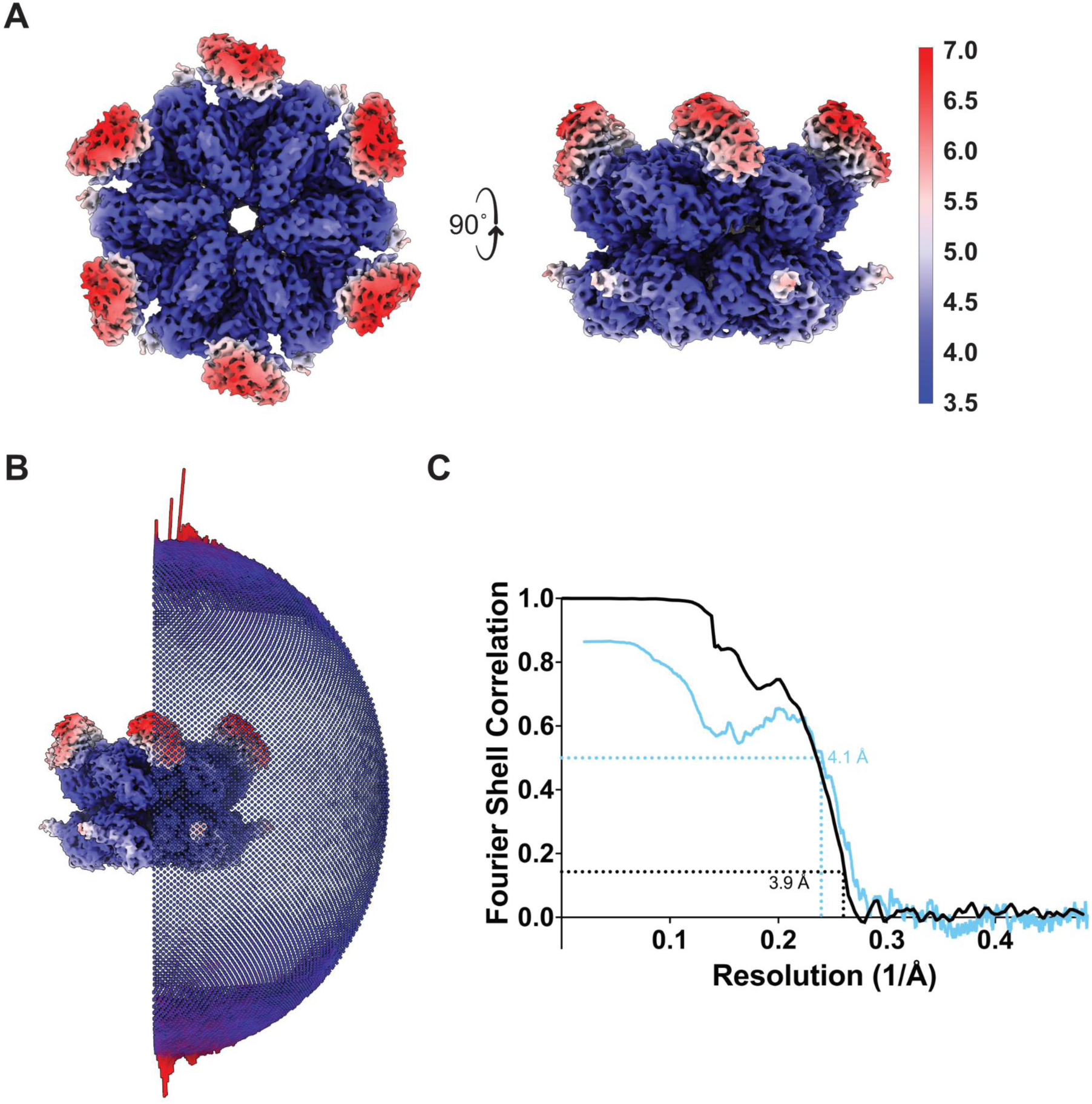
Cryo-EM reconstruction validation metrics of apo AtCDC48A. **A)** Cryo-EM density coloured by local resolution. Low–high resolution regions are coloured red–blue. **B)** Euler angle distribution of the final reconstruction with C6 symmetry. **C)** Fourier shell correlation (FSC) curves for the cryo-EM reconstructions. Black line represents the correlation between independently process half maps, with resolution determined at FSC = 0.143. Blue line shows the correlation between the atomic model (**PDB 9M3V**) and the cryo-EM reconstruction (**EMD-63608**), with the resolution assessed at FSC = 0.5.

**Figure S4:**
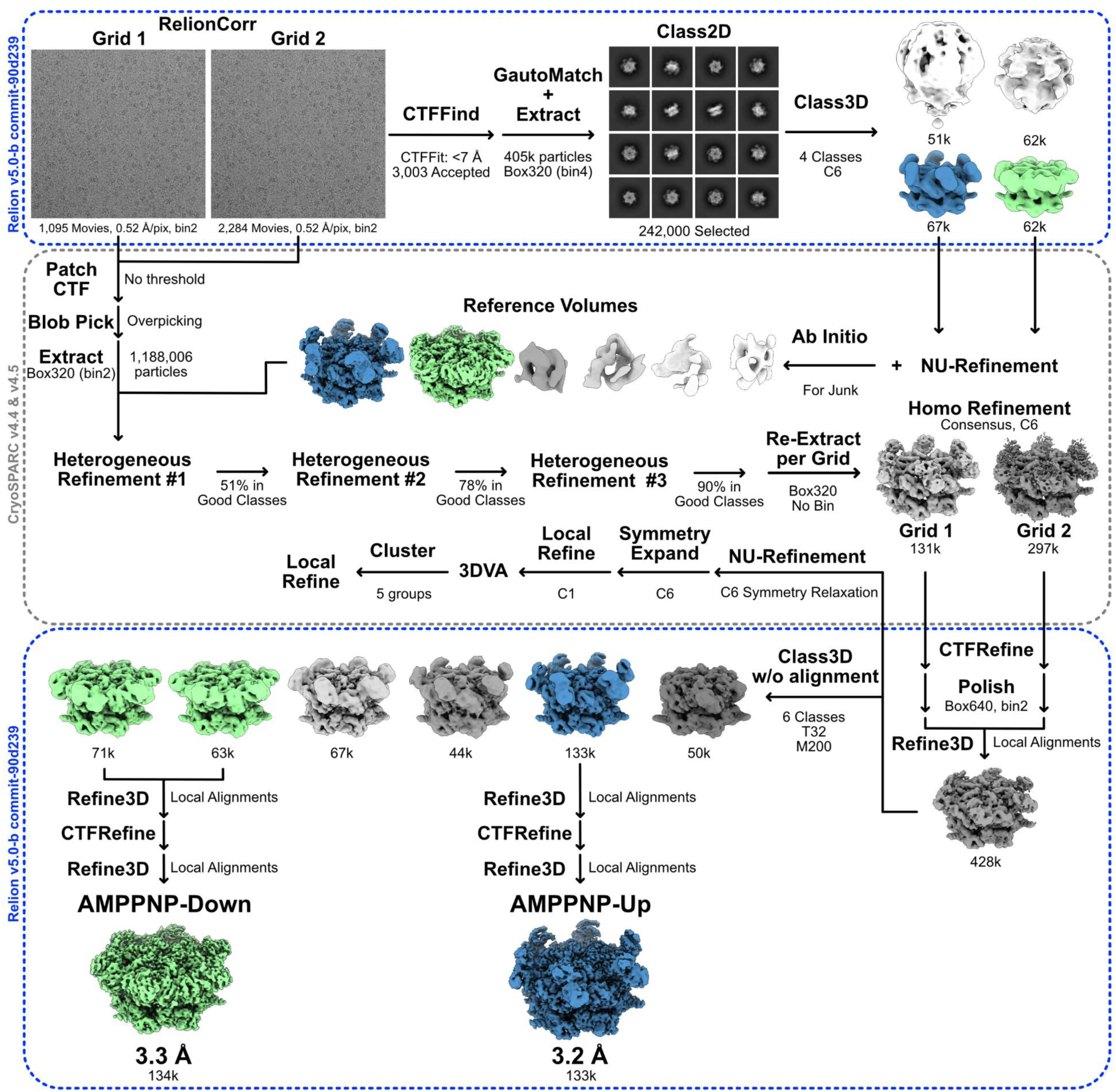
Cryo-EM image processing of AMPPNP-bound *At*CDC48A.

**Figure S5:**
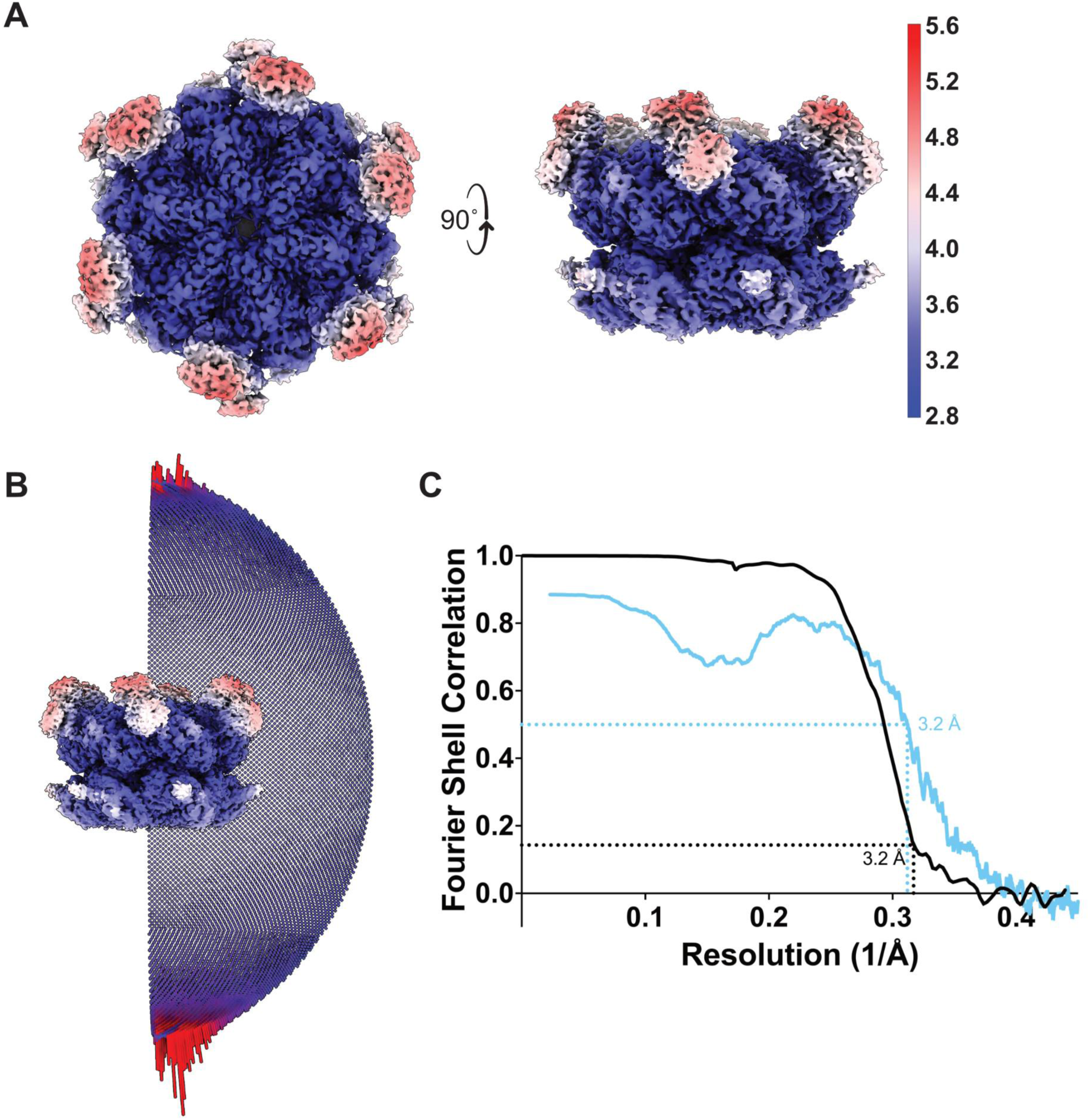
Cryo-EM reconstruction validation metrics of *At*CDC48A in AMPPNP- Up state. **A)** Cryo-EM density coloured by local resolution. Low–high resolution regions are coloured red–blue. **B)** Euler angle distribution of the final reconstruction with C6 symmetry. **C)** Fourier shell correlation (FSC) curves for the cryo-EM reconstructions. Black line represents the correlation between independently process half maps, with resolution determined at FSC = 0.143. Blue line shows the correlation between the atomic model (**PDB 9M3W**) and the cryo-EM reconstruction (**EMD-63609**), with the resolution assessed at FSC = 0.5.

**Figure S6:**
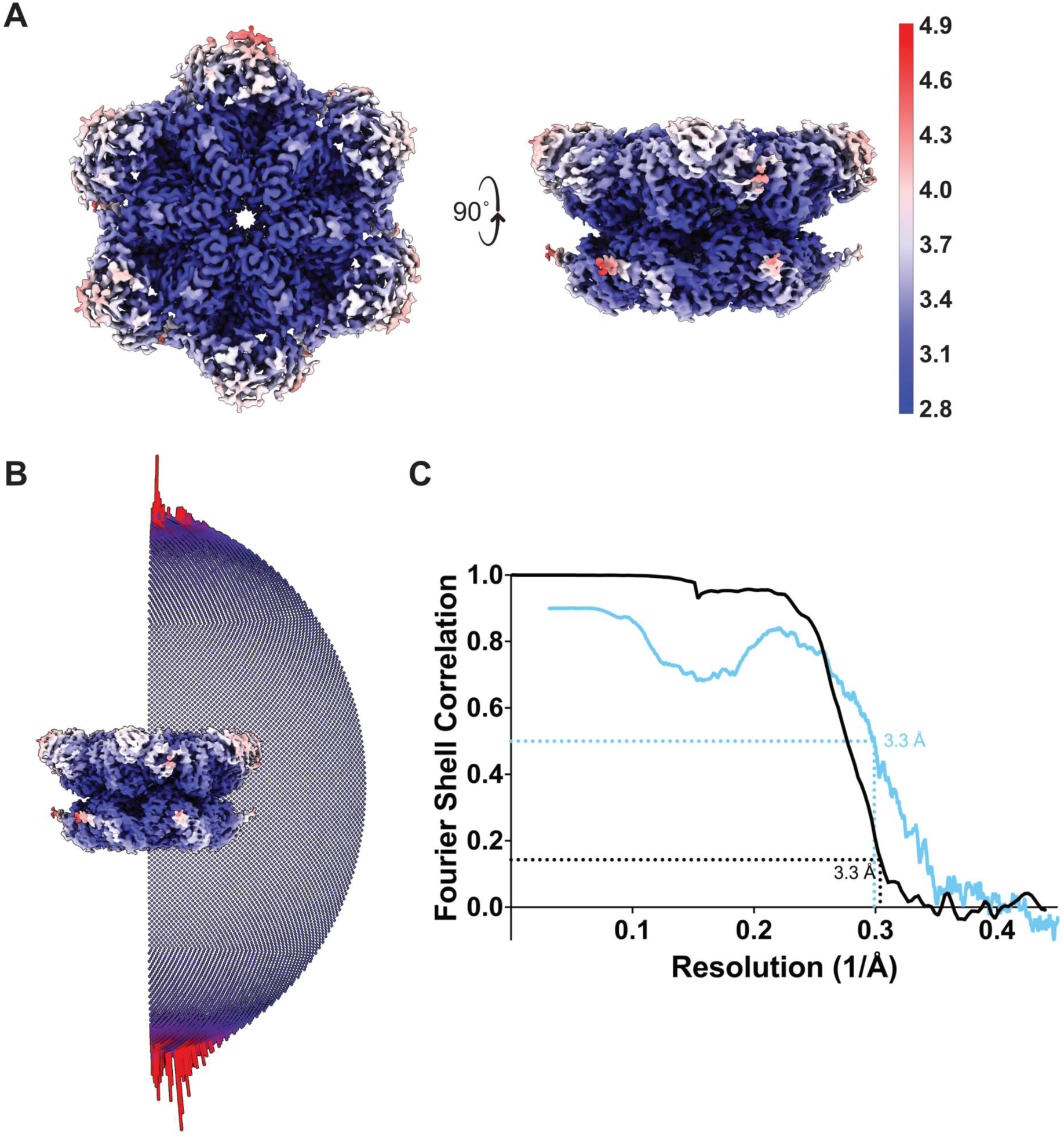
Cryo-EM reconstruction validation metrics of *At*CDC48A in AMPPNP- Down state. **A)** Cryo-EM density coloured by local resolution. Low–high resolution regions are coloured red–blue. **B)** Euler angle distribution of the final reconstruction with C6 symmetry. **C)** Fourier shell correlation (FSC) curves for the cryo-EM reconstructions. Black line represents the correlation between independently process half maps, with resolution determined at FSC = 0.143. Blue line shows the correlation between the atomic model (**PDB 9M3X**) and the cryo-EM reconstruction (**EMD-63610**), with the resolution assessed at FSC = 0.5.

**Figure S7:**
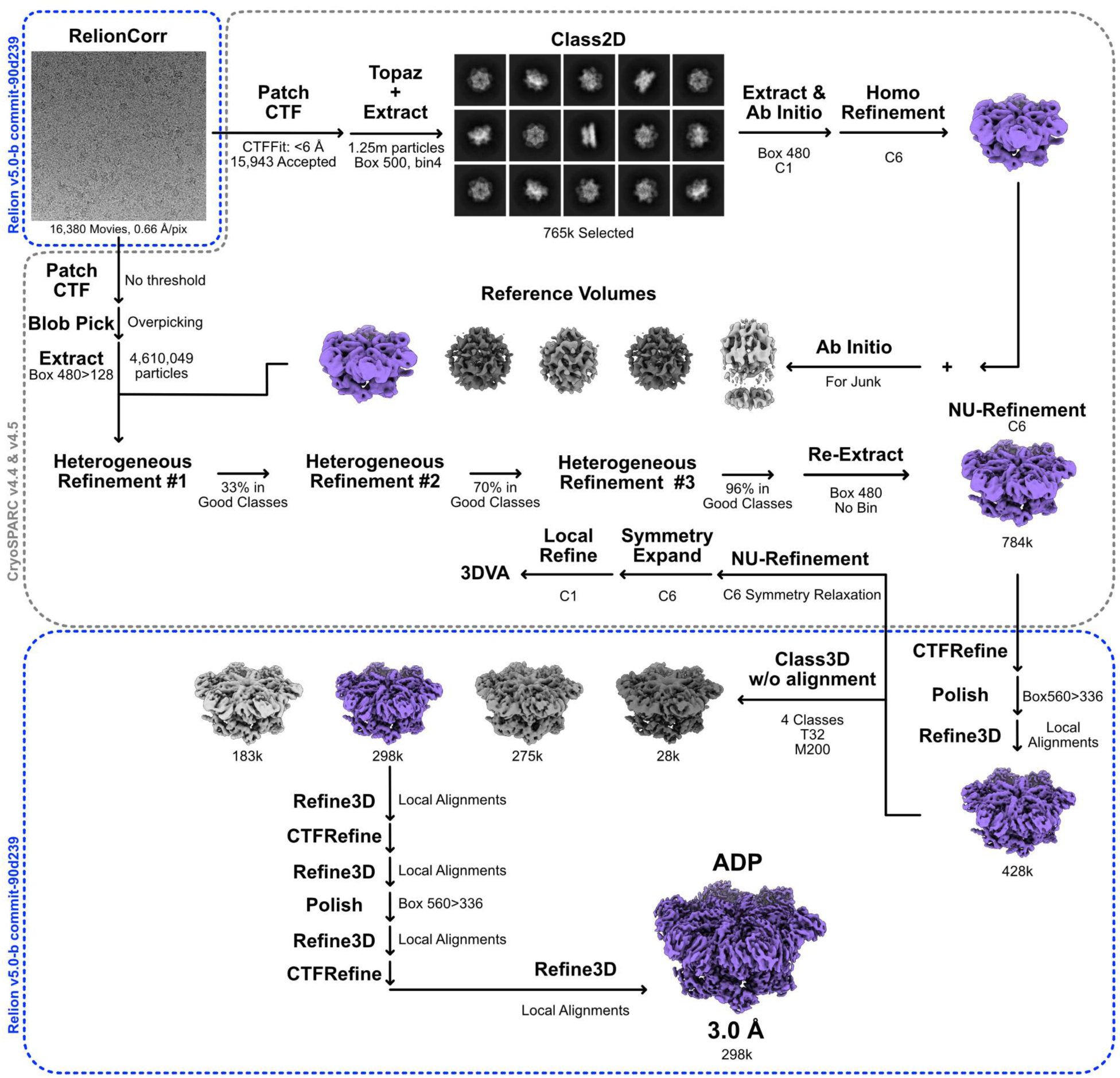
Cryo-EM image processing of ADP-bound *At*CDC48A.

**Figure S8:**
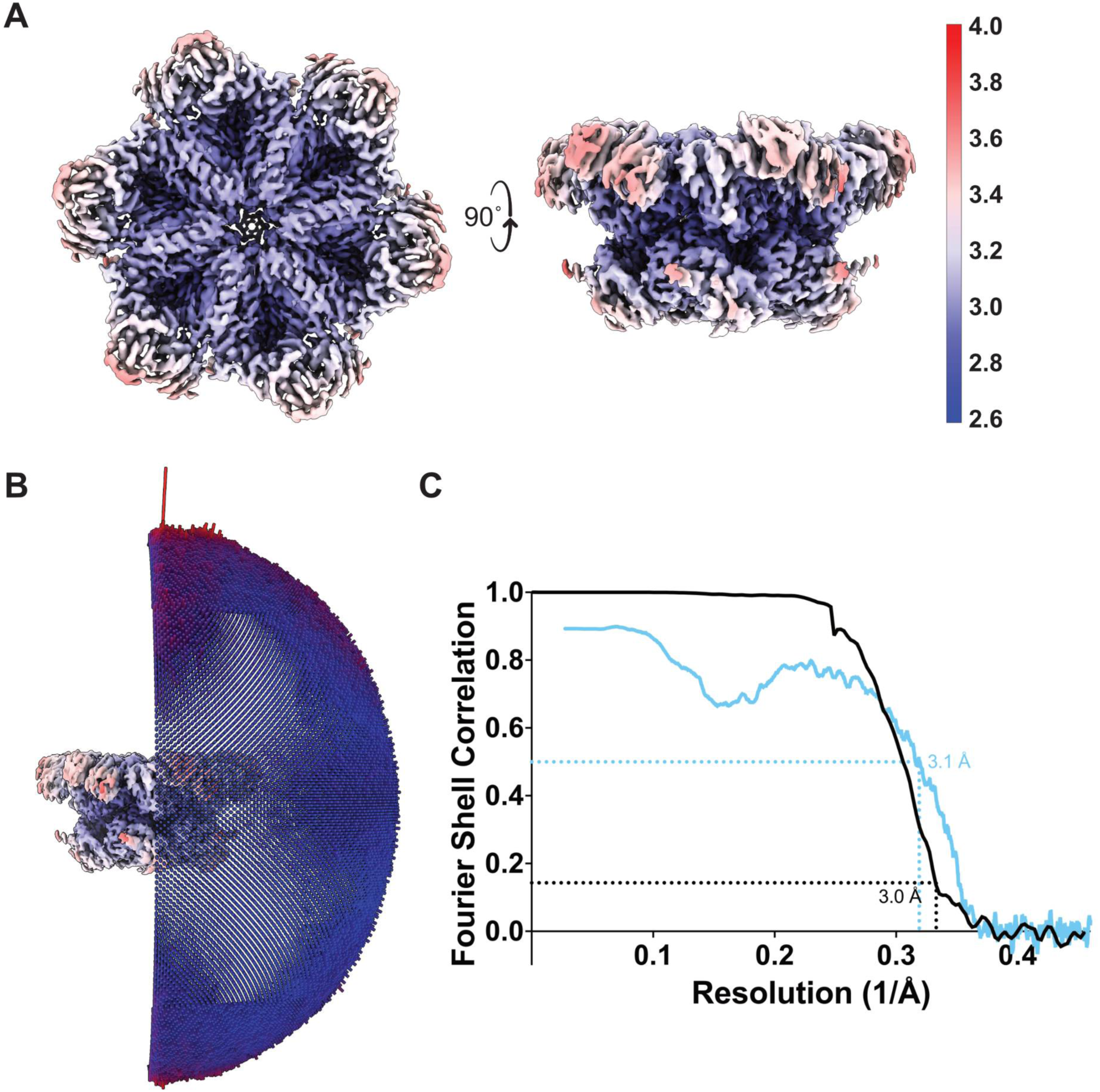
Cryo-EM reconstruction validation metrics of ADP-bound *At*CDC48A. **A)** Cryo-EM density coloured by local resolution. Low–high resolution regions are coloured red–blue. **B)** Euler angle distribution of the final reconstruction with C6 symmetry. **C)** Fourier shell correlation (FSC) curves for the cryo-EM reconstructions. Black line represents the correlation between independently process half maps, with resolution determined at FSC = 0.143. Blue line shows the correlation between the atomic model (**PDB 9M3Y**) and the cryo-EM reconstruction (**EMD-63611**), with the resolution assessed at FSC = 0.5.

**Figure S9:**
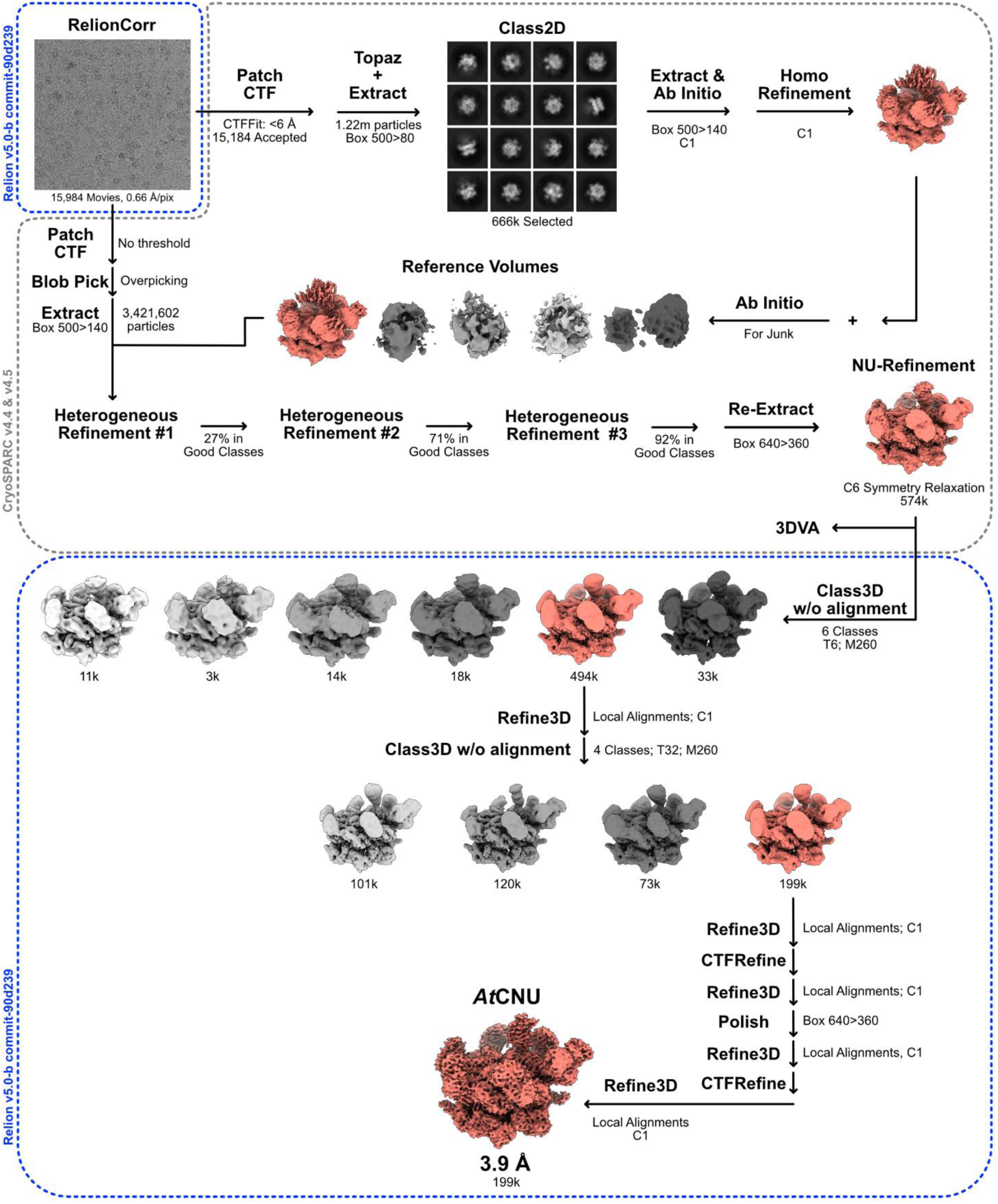
Cryo-EM image processing of *At*CNU.

**Figure S10:**
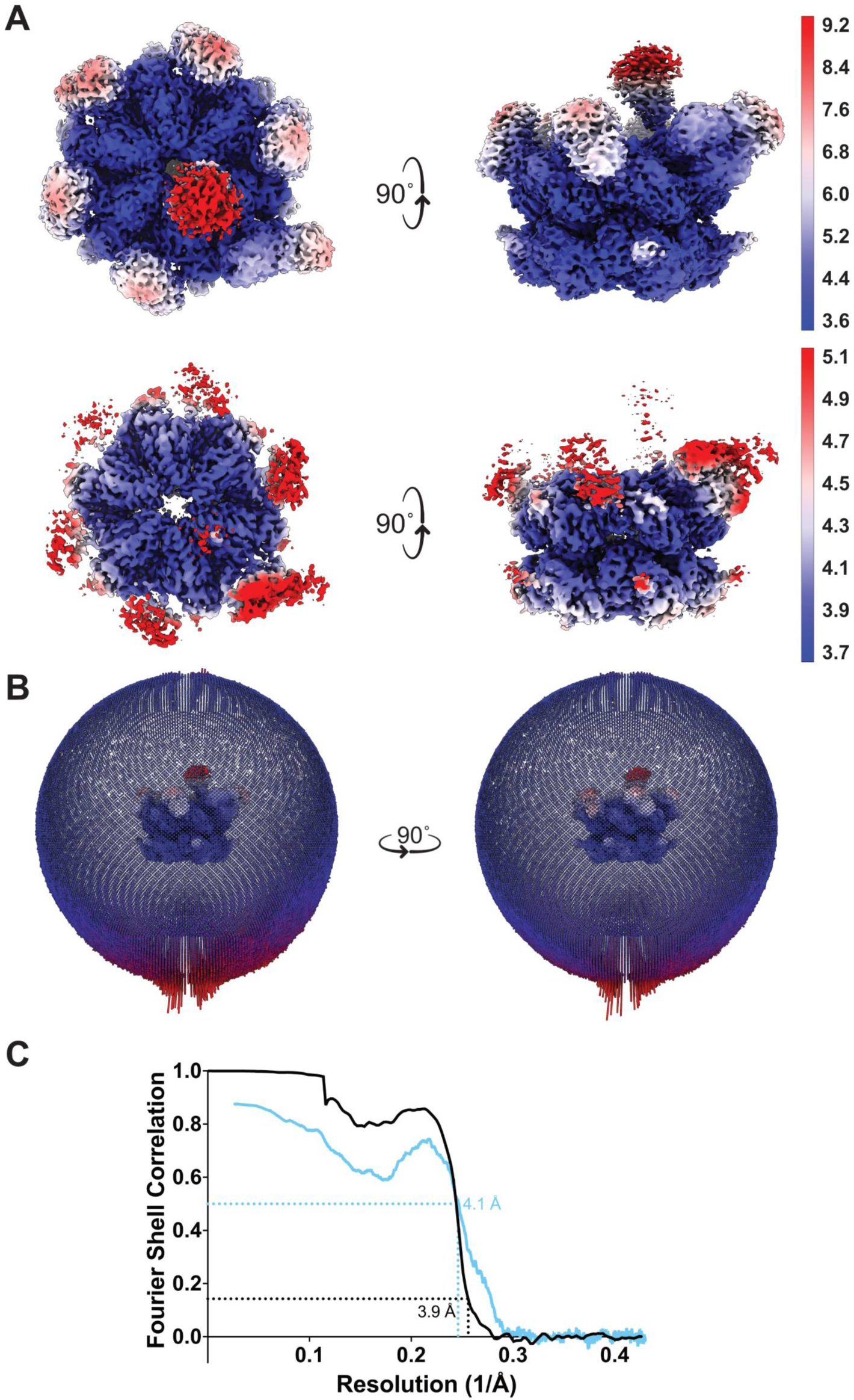
Cryo-EM reconstruction validation metrics of *At*CNU. **A)** Cryo-EM density coloured by local resolution. Low–high resolution regions are coloured red–blue. **B)** Euler angle distribution of the final reconstruction with C6 symmetry. **C)** Fourier shell correlation (FSC) curves for the cryo-EM reconstructions. Black line represents the correlation between independently process half maps, with resolution determined at FSC = 0.143. Blue line shows the correlation between the atomic model (**PDB 9M3Z**) and the cryo-EM reconstruction (**EMD-63612**), with the resolution assessed at FSC = 0.5.

**Figure S11.**
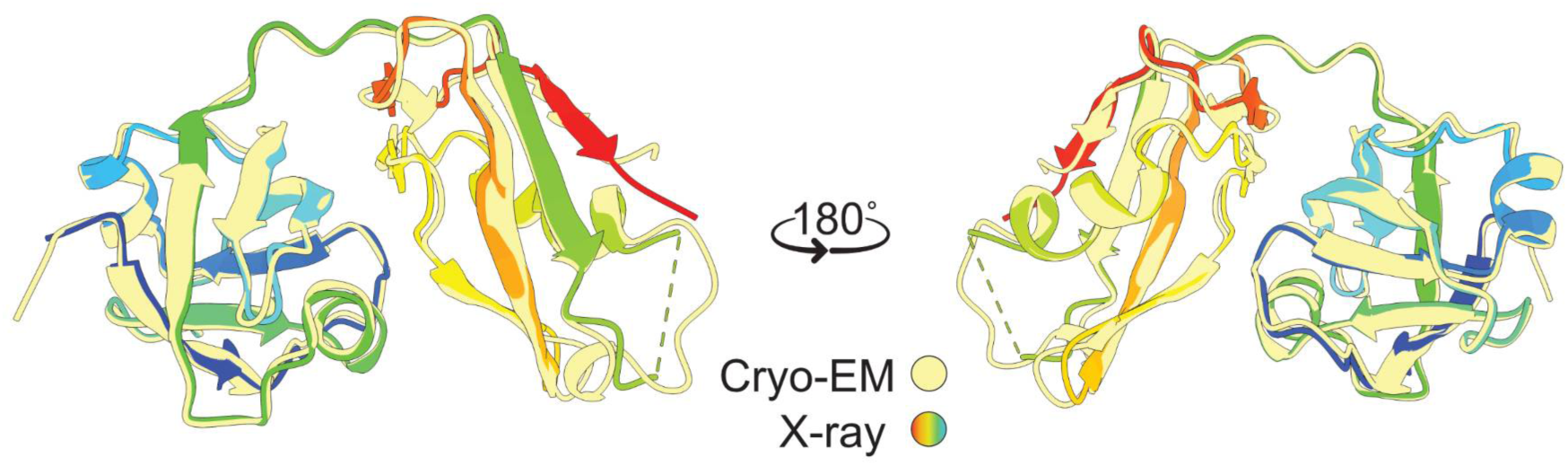
Crystal structure of the *At*CDC48A N-terminal Domain (NTD). The NTD of the ADP-bound cryo-EM model (**PDB 9M3Y**; **EMD 63611**) is coloured yellow, and the NTD crystal structure (**PDB 9M4G**) is coloured as a rainbow (blue–red) by sequence. Dashed lines indicate unmodelled residues.

**Figure S12.**
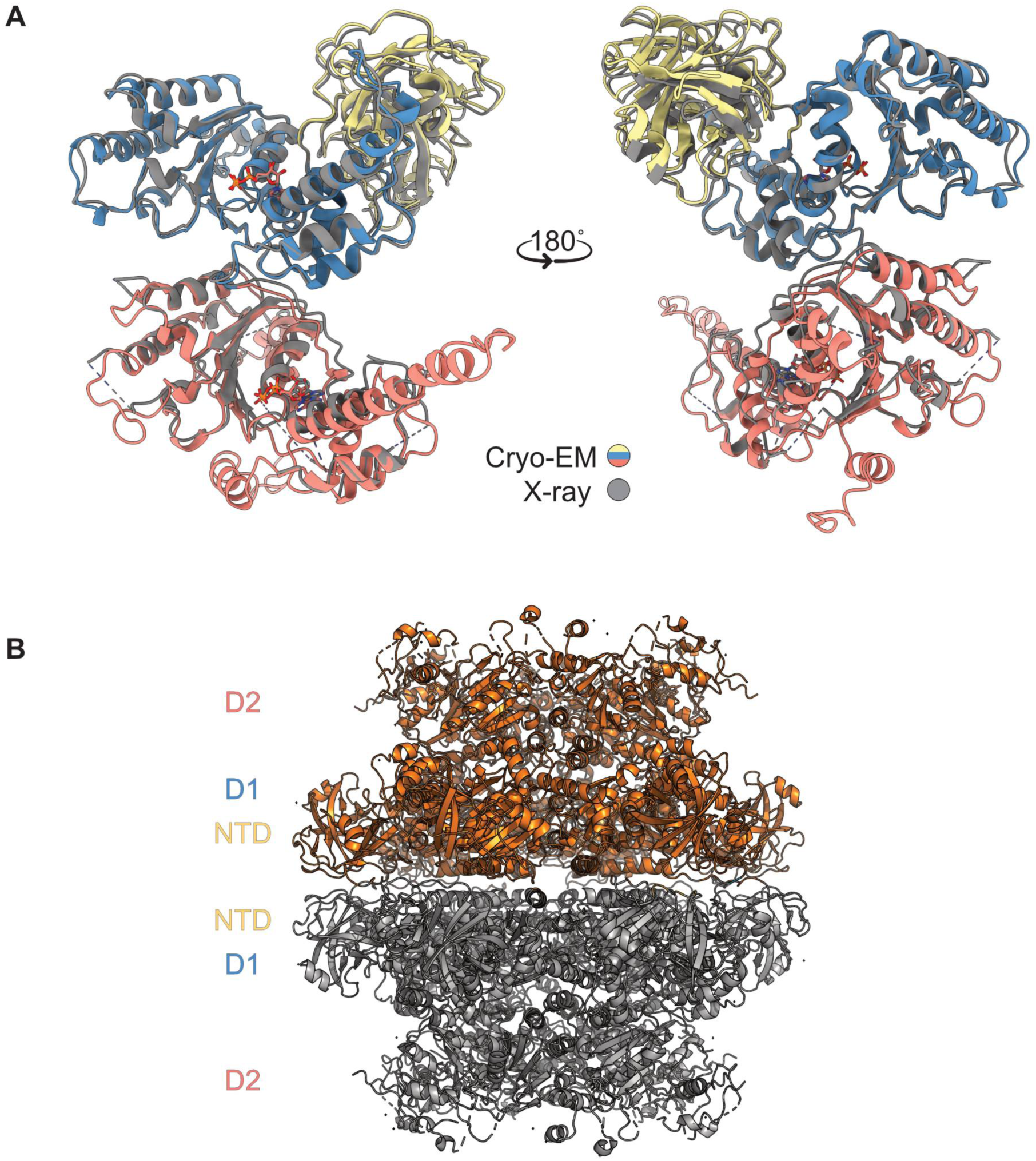
Crystal structure of ADP-bound *At*CDC48A_28-809_. **A)** One protomer of the crystal structure is superimposed on the cryo-EM structure. The ADP-bound cryo-EM structure (**PDB 9M3Y**; **EMD 63611**) is coloured as in Figure 1. The crystal structure (**PDB 9M4N**) is coloured grey. ADP molecules are depicted as sticks. Dashed lines indicate unmodelled residues. **B)** Dodecameric arrangement of *At*CDC48A_28-809_. One hexamer is coloured orange and the other coloured grey. The NTD, D1, and D2 position is labelled.

**Figure S13.**
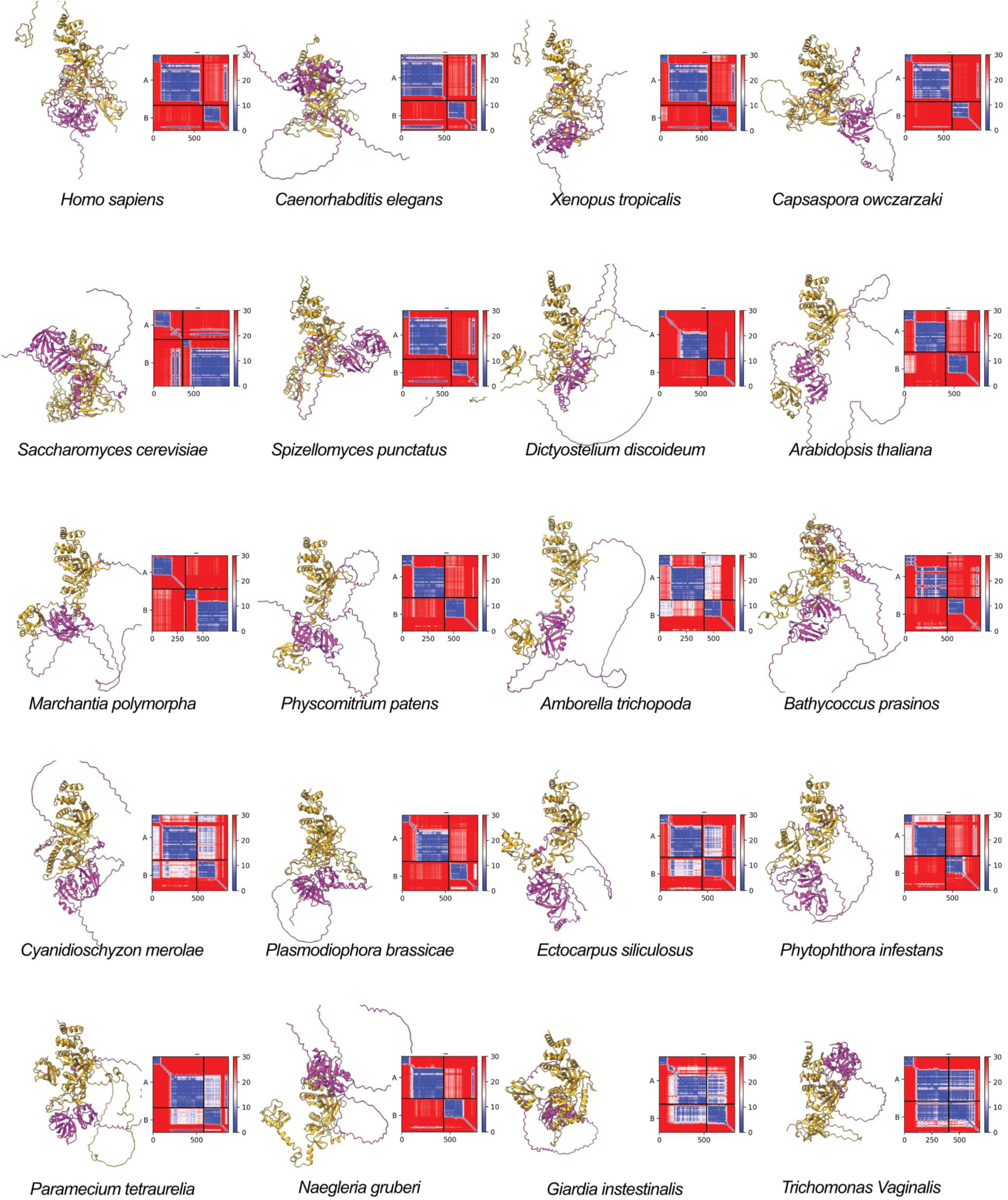
AlphaFold predicted models of NPL4–UFD1 across species. For each species, the predicted model (left) and PAE plot (left) are depicted, shown with scale bar representing low (blue) to high (red) PAE. NPL4 and UFD1 are coloured orange and pink, respectively.

**Figure S14.**
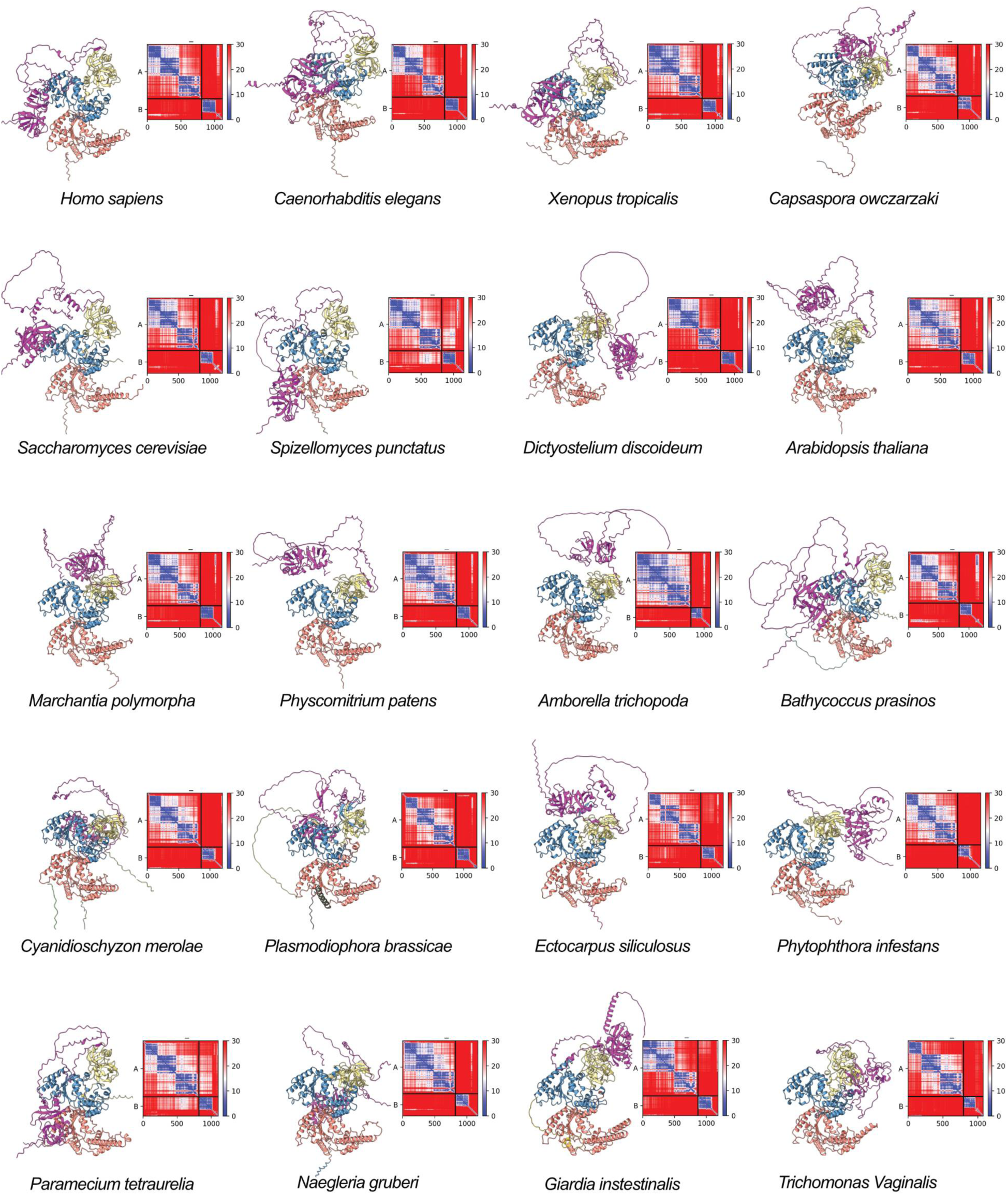
AlphaFold predicted models of CDC48–UFD1 across species. For each species, the predicted model (left) and PAE plot (left) are depicted, shown with scale bar representing low (blue) to high (red) PAE. CDC48 is coloured by domain and UFD1 is coloured pink.

**Figure S15.**
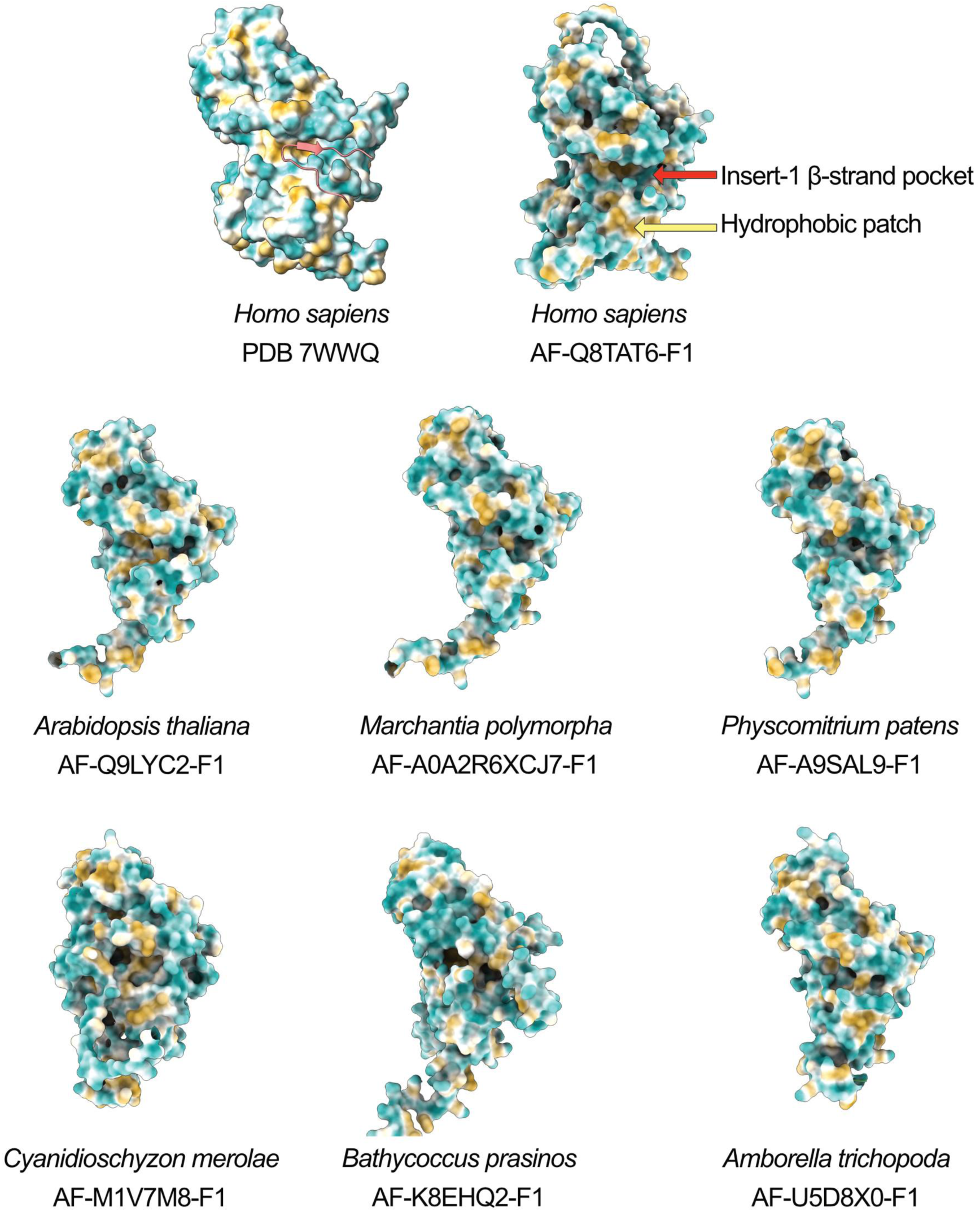
Analysis of NPL4’s UFD1-binding region (UBR). The crystal structure of the *Hs*Npl4–*Hs*Ufd1 heterodimer (PDB 7WWQ) is shown on the top left for reference. The AlphaFold predicted structure of *Hs*Npl4’s MPN domain is shown on the top right, with the insert-1 B-strand pocket (red arrow) and hydrophobic patch (yellow arrow) labelled. Below, AlphaFold predicted structures of representative plant species’ NPL4 MPN domains. The AlphaFold directory accession code is provided for each model. The surface of the MPN domain is depicted and coloured blue–gold for hydrophilic– hydrophobic residues. Residues 1-100 are hidden for clarity.

**Figure S16.**
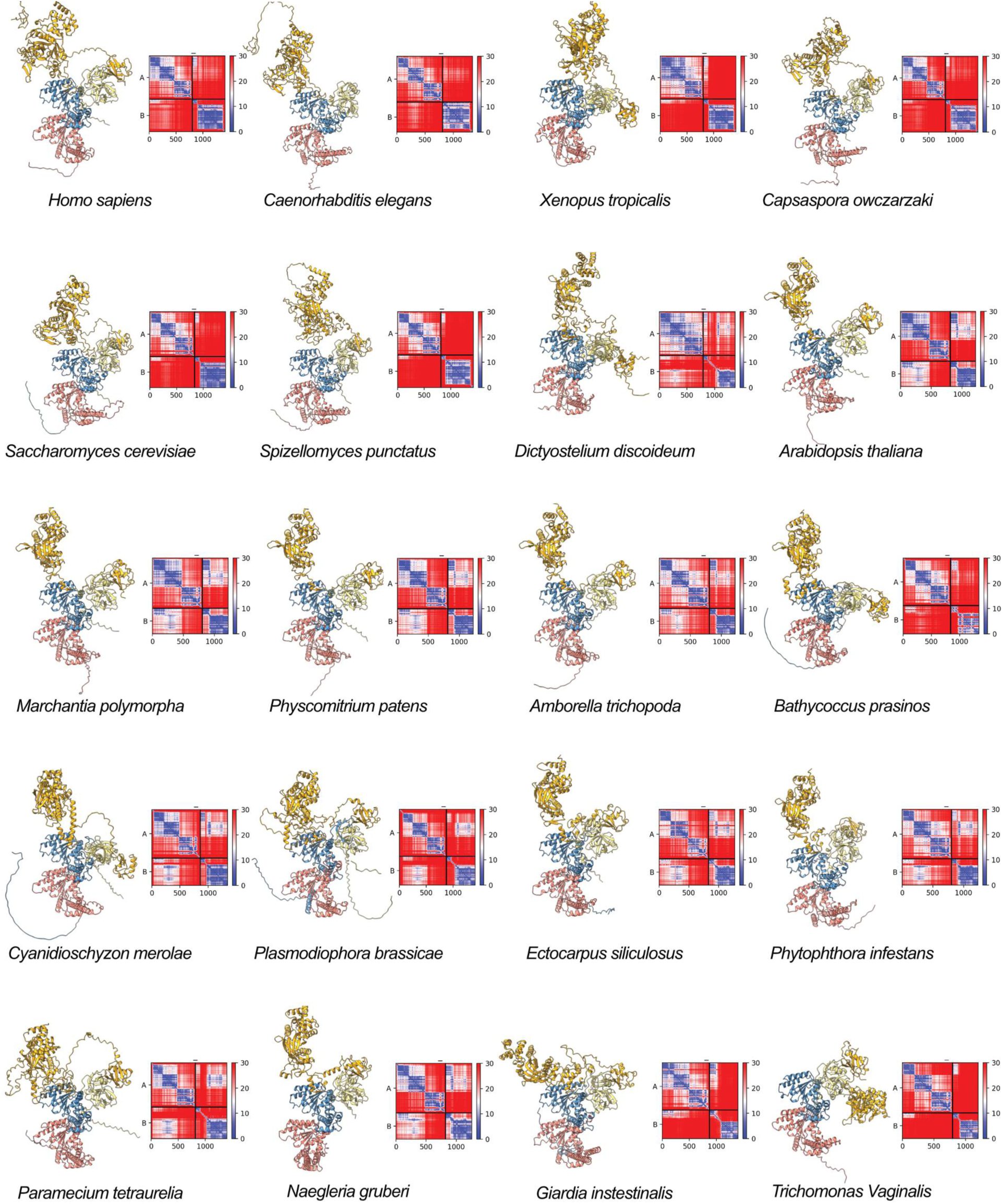
AlphaFold predicted models of CDC48–NPL4 across species. For each species, the predicted model (left) and PAE plot (left) are depicted, shown with scale bar representing low (blue) to high (red) PAE. CDC48 is coloured by domain and NPL4 is coloured orange.

**Figure S17.**
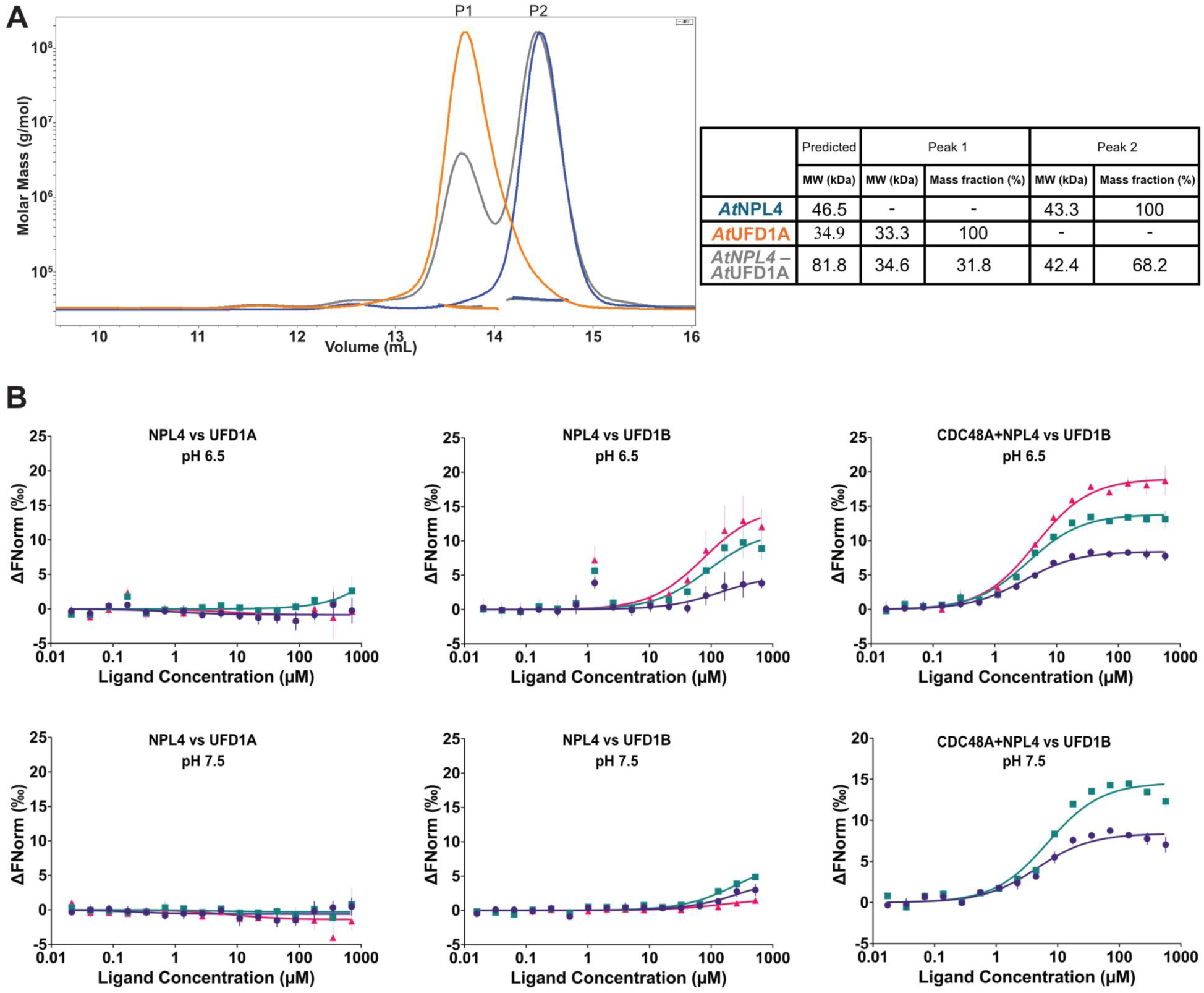
Interaction of the *At*CNU complex. **A)** *Left panel:* Differential refractive index (solid curves) of *At*CNU proteins measured through Size Exclusion Chromatography coupled with Multi-Angle Light Scattering (SEC-MALS). The calculated molecular weight of each peak (P1–P2) is represented by a dashed line and tabulated on the right. *Right panel:* Tabulated molecular weights calculated for each individual protein or complex. The predicted molecular weight of each species is provided, alongside the calculated values for each peak on the left (P1–P2). *At*NPL4 is coloured blue, *At*UFD1A is orange and *At*NPL4–*At*UFD1A is grey. **B)** Representative microscale thermophoresis (MST) interaction plots at pH 6.5 (top) and pH 7.5 (bottom) for *At*NPL4– *At*UFD1A (left), *At*NPL4–*At*UFD1B (middle) and *At*CDC48A saturated with *At*NPL4 to *At*UFD1B (right). The change of normalised fluorescence (y-axis) is plotted as a function of ligand (*At*UFD1A or *At*UFD1B) concentration. Purple, teal, and magenta points and lines of best fit represent triplicate measurements at low (20%), medium (40%) and high (60%) MST power, respectively.

**Figure S18.**
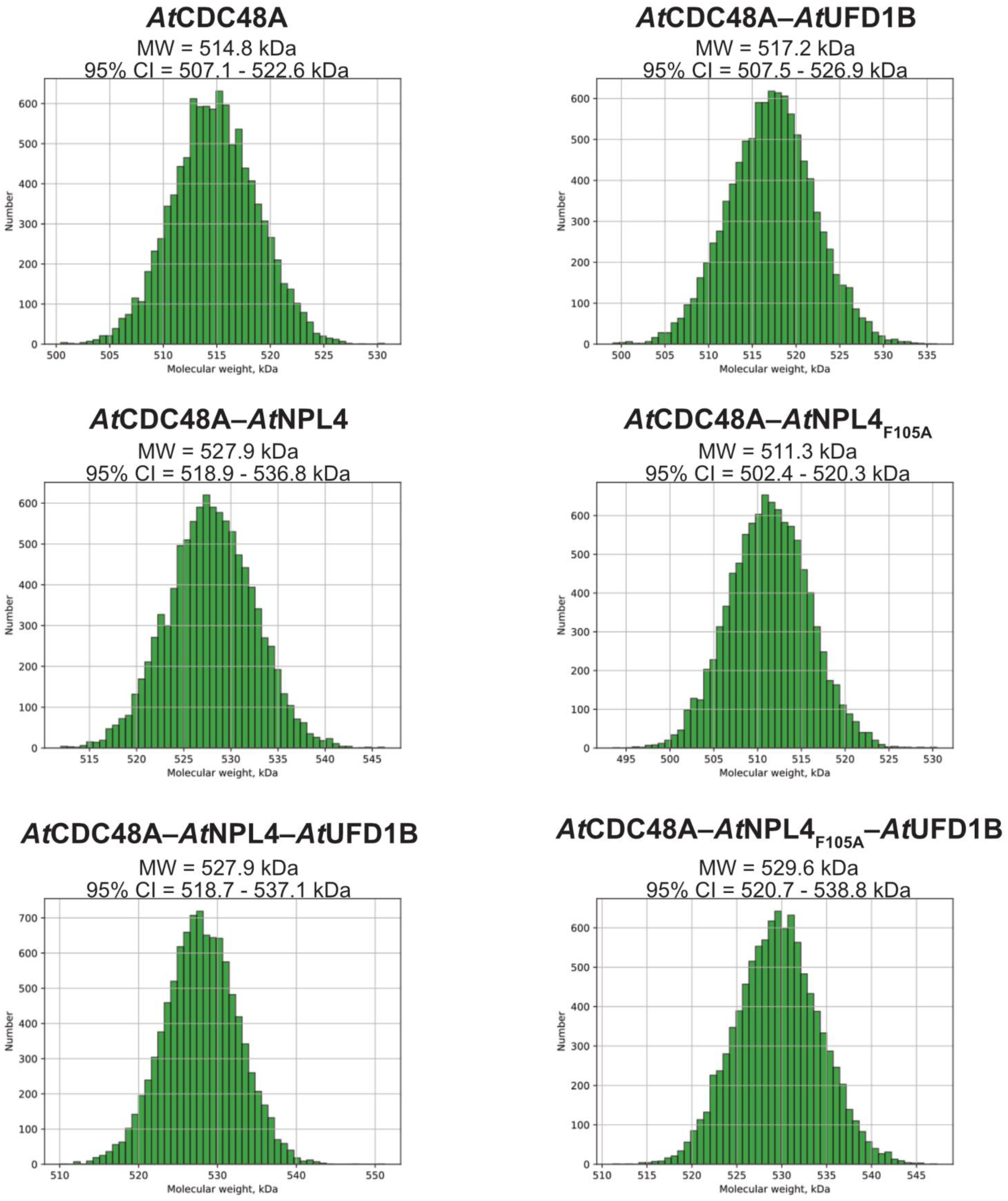
Molecular weight of *At*CNU complexes by SEC-SAXS. The complex and its molecular weight (MW) and 95% confidence intervals (CI) are provided above the corresponding histogram. Molecular weights were calculated from the one-dimensional scattering curve using the MW-DARA web server.

**Figure S19.**
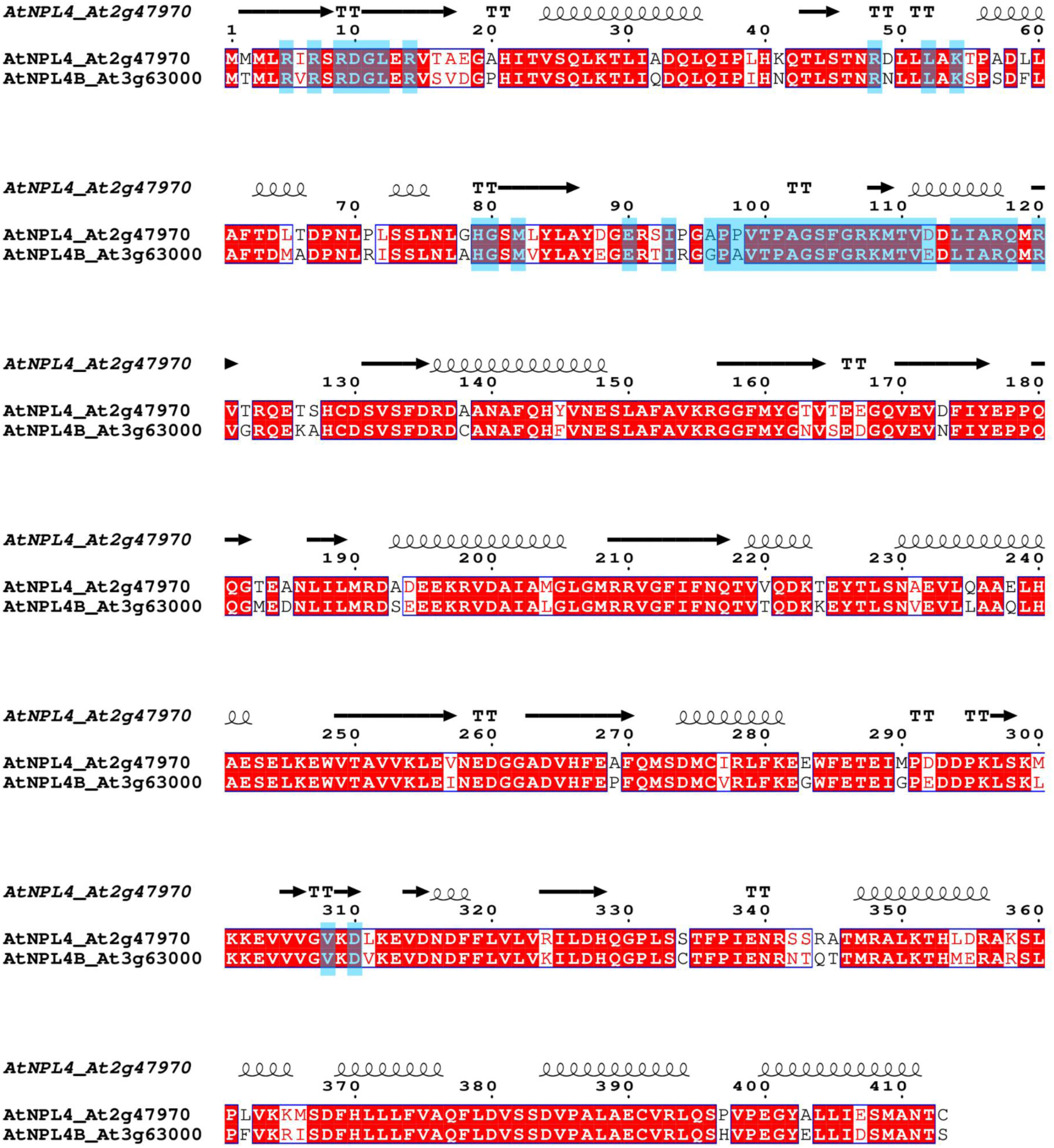
Sequence alignment of *At*NPL4 and *At*NPL4B. Secondary structure features and residue numbers are provided for AtNpl4 above the alignment. Red colouring indicates identical residues. AtCDC48A-interacting residues from the AtCNU cryo-EM model are highlighted blue.

**Figure S20.**
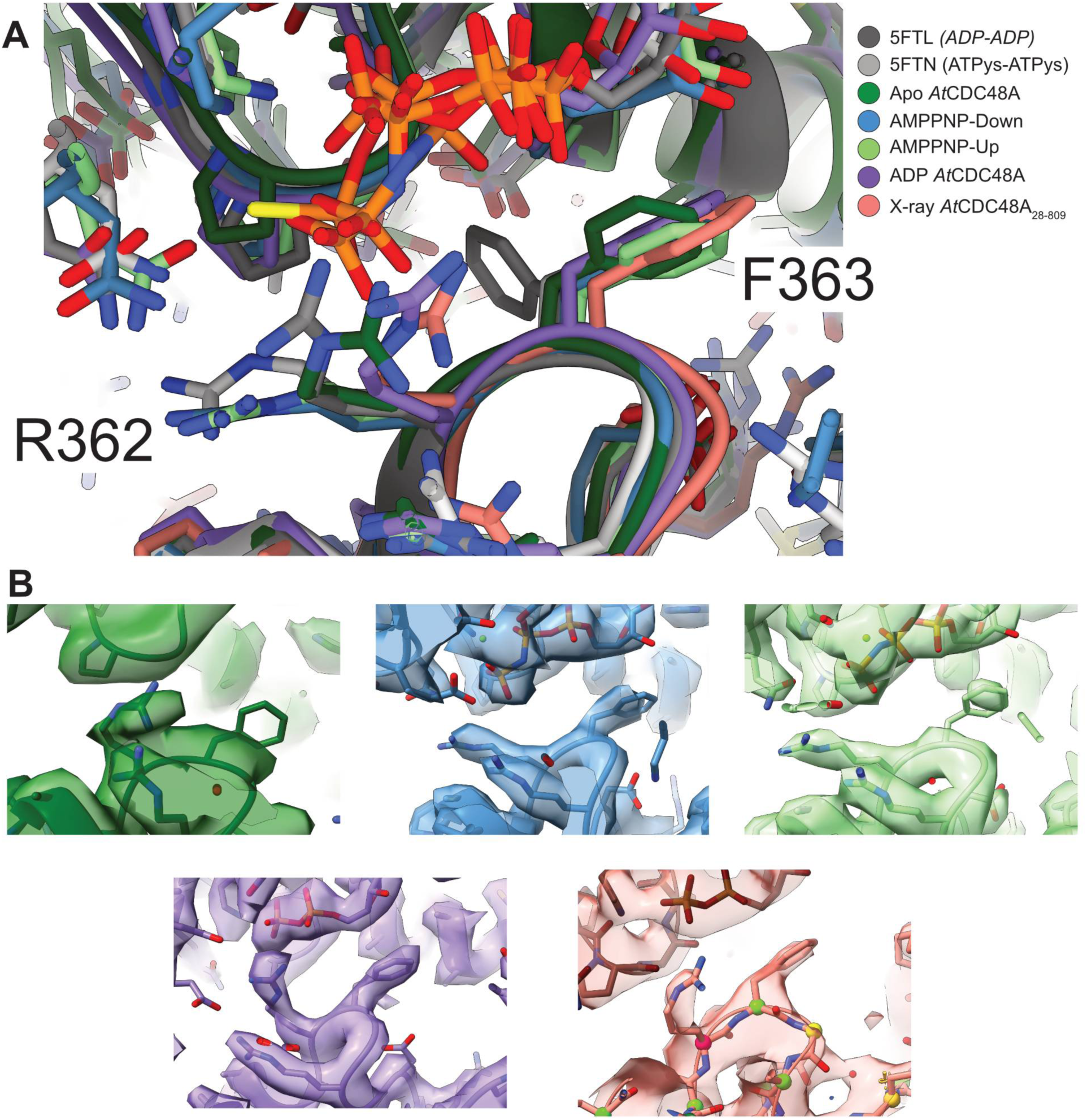
Comparison of F360 across *At*CDC48A structures. **A)** Superimposition of D1 atomic models from *Hs*p97 in ADP (dark grey; **PDB 5FTL**) and ATPᵧS (light grey; **PDB 5FTL**) states and *At*CDC48A cryo-EM models of the apo-state (dark green; **PDB 9M3V**), AMPPNP-Up (blue; **PDB 9M3W**), AMPPNP-Down (green; **PDB 9M3X**) and ADP-state (purple; **PDB 9M3Y**) and the ADP-bound X-ray model (red; **PDB 9M4G**). Atoms are represented as sticks. Residues R362 and F363 are labelled. **B)** Local fit of models to their respective EM or electron density map for each of the *At*CDC48A models in Panel A. Coloured as in Panel A.

**Table S1:** Cryo-EM data collection, refinement and validation statistics.

|  | **Apo** | **AMPPNP-Up** | **AMPPNP-Down** | **ADP** | **AtCNU** |
| --- | --- | --- | --- | --- | --- |
|  | EMD-63608 | EMD-63609 | EMD-63610 | EMD-63611 | EMD-63612 |
|  | PDB 9M3V | PDB 9M3W | PDB 9M3X | PDB 9M3Y | PDB 9M3Z |
| **Data collection and processing** |  |  |  |  |  |
| Magnification | 130k | 130k | 130k | 130k | 130k |
| Voltage (kV) | 300 | 300 | 300 | 300 | 300 |
| Electron exposure (e-/Å<sup>2</sup>) | 50 | 50 | 50 | 64 | 66 |
| Defocus range (μm) | -2.2 – -1.0 | -2.2 – -1.0 | -2.2 – -1.0 | -2.6 – -0.8 | -2.6 – -0.8 |
| Pixel size (Å) | 0.52 | 0.52 | 0.52 | 0.66 | 0.66 |
| Symmetry imposed | C6 | C6 | C6 | C6 | C1 |
| Initial particle images (no.) | 201k | 1.2m | 1.2m | 4.6m | 3.4m |
| Final particle images (no.) | 73k | 133k | 134k | 298k | 199k |
| Map resolution (Å) |  |  |  |  |  |
| (FSC threshold 0.143) | 3.9 | 3.2 | 3.3 | 3.0 | 3.9 |
| Map resolution range (Å) | 3.7 – 7.9 | 3.0 – 5.6 | 3.1 – 5.2 | 2.8 – 4.3 | 3.7 – 10.4 |
| **Refinement** |  |  |  |  |  |
| Initial model used (PDB code) | AlphaFold | AlphaFold | AlphaFold | AlphaFold | AlphaFold |
| Model resolution (Å) |  |  |  |  |  |
| (FSC threshold 0.5) | 4.1 | 3.2 | 3.3 | 3.1 | 4.1 |
| Map sharpening *B* factor (Å<sup>2</sup>) | -127.3 | -80.4 | -100.3 | -100.0 | -80.4 |
| Model composition |  |  |  |  |  |
| Non-hydrogen atoms | 35,280 | 36,012 | 35,502 | 35,700 | 39,251 |
| Protein residues | 753 | 4566 | 4506 | 4530 | 4979 |
| Ligands | 0 | 24 | 24 | 12 | 18 |
| *B* factors (Å<sup>2</sup>) |  |  |  |  |  |
| Protein | 55.52 | 70.19 | 56.56 | 64.19 | 39.32 |
| Ligand | - | 15.73 | 21.01 | 19.45 | 21.15 |
| R.m.s. deviations |  |  |  |  |  |
| Bond lengths (Å) | 0.003 | 0.003 | 0.003 | 0.002 | 0.002 |
| Bond angles (°) | 0.548 | 0.484 | 0.449 | 0.557 | 0.519 |
| Validation |  |  |  |  |  |
| MolProbity score | 1.67 | 1.61 | 1.27 | 1.86 | 2.07 |
| Clashscore | 8.66 | 4.55 | 2.73 | 5.83 | 8.88 |
| Poor rotamers (%) | 0.16 | 0.78 | 0.79 | 2.34 | 2.20 |
| Ramachandran plot |  |  |  |  |  |
| Favoured (%) | 96.78 | 94.33 | 96.66 | 96.15 | 95.25 |
| Allowed (%) | 3.22 | 5.53 | 3.07 | 3.59 | 4.73 |
| Disallowed (%) | 0.00 | 0.13 | 0.27 | 0.27 | 0.02 |

**Table S2:** X-ray crystallography statistics.

|  | **AtCDC48A<sub>28-809</sub>**<br>PDB 9M4N | **NTD (AtCDC48A<sub>28-190</sub>)**<br>PDB 9M4G |
| --- | --- | --- |
| Wavelength | 0.980118 | 0.980112 |
| Resolution range | 47.29–3.45 | 48.25–2.60 |
| Space group | P 6 2 2 | R 3 2 :H |
| Unit cell | 144.47 144.47 162.96 90 90<br>120 | 91.72 91.72 242.98 90 90<br>120 |
| Total reflections | 439430 (104460) | 253189 (31188) |
| Unique reflections | 13805 (3211) | 12475 (1486) |
| Multiplicity | 31.8 (32.5) | 20.3 (21.0) |
| Completeness (%) | 100 (100.00) | 100 (100.00) |
| Mean I/sigma(I) | 8.4 (1.5) | 10.4 (1.2) |
| Wilson B-factor | 85.93 | 83.1 |
| R-merge | 0.62 (3.77) | 0.199 (3.143) |
| R-meas | 0.64 (3.88) | 0.209 (3.297) |
| R-pim | 0.15 (0.92) | 0.064 (0.996) |
| CC1/2 | 0.99 (0.32) | 0.998 (0.308) |
| Reflections used in refinement | 13799 (1341) | 12455 (1235) |
| Reflections used for R-free | 701 (68) | 657 (80) |
| R-work | 0.3461 (0.4567) | 0.2438 (0.3770) |
| R-free | 0.3838 (0.4478) | 0.3189 (0.4247) |
| Number of non-hydrogen atoms | 5207 | 2521 |
| macromolecules | 5143 | 2502 |
| ligands | 55 | 5 |
| solvent | 9 | 14 |
| Protein residues | 659 | 314 |
| RMS(bonds) | 0.003 | 0.011 |
| RMS(angles) | 0.68 | 1.36 |
| Ramachandran favored (%) | 88.8 | 91.12 |
| Ramachandran allowed (%) | 11.04 | 8.88 |
| Ramachandran outliers (%) | 0.16 | 0 |
| Rotamer outliers (%) | 0.53 | 0.71 |
| Clashscore | 11.98 | 16.23 |
| Average B-factor | 100.98 | 98.71 |
| macromolecules | 100.7 | 98.78 |
| ligands | 129.76 | 105.23 |
| solvent | 84.89 | 82.72 |

**Table S3: Intermolecular crosslinking mass spectrometry peptides between *At*CDC48A–*At*UFD1B.** Complete list of intermolecular crosslinks between *At*CDC48A and *At*UFD1B. Complexes were crosslinked using the linker BS3. Peptides including the SHP1 motif are highlighted light green. Peptides surrounding the SHP1 motif are highlighted dark green. Peptides including the SHP2 motif are coloured orange.

**Table S4: Intermolecular crosslinking mass spectrometry peptides between *At*CDC48A–*At*NPL4.** Complete list of intermolecular crosslinks between *At*CDC48A and *At*NPL4. Complexes were crosslinked using the linker BS3. Peptides including the UBXL domain are coloured dark blue. Peptides including the UBXL–MPN linker are coloured light blue.

**Table S5: Sequences and accession numbers for phylogenetic analysis.**

